# A tutorial on Gaussian process regression: Modelling, exploring, and exploiting functions

**DOI:** 10.1101/095190

**Authors:** Eric Schulz, Maarten Speekenbrink, Andreas Krause

**Affiliations:** Department of Psychology, Harvard University; Department of Experimental Psychology, University College London; Department of Computer Science, Swiss Federal Institute of Technology Zürich

**Keywords:** Gaussian Process Regression, Active Learning, Exploration-Exploitation, Bandit Problems

## Abstract

This tutorial introduces the reader to Gaussian process regression as an expressive tool to model, actively explore and exploit unknown functions. Gaussian process regression is a powerful, non-parametric Bayesian approach towards regression problems that can be utilized in exploration and exploitation scenarios. This tutorial aims to provide an accessible introduction to these techniques. We will introduce Gaussian processes which generate distributions over functions used for Bayesian non-parametric regression, and demonstrate their use in applications and didactic examples including simple regression problems, a demonstration of kernel-encoded prior assumptions and compositions, a pure exploration scenario within an optimal design framework, and a bandit-like exploration-exploitation scenario where the goal is to recommend movies. Beyond that, we describe a situation modelling risk-averse exploration in which an additional constraint (not to sample below a certain threshold) needs to be accounted for. Lastly, we summarize recent psychological experiments utilizing Gaussian processes. Software and literature pointers are also provided.

## 1. Introduction

Whether we try to find a function that accurately describes participants’ behaviour (Cavagnaro, Aranovich, McClure, Pitt, and Myung, 2014), estimate parameters of psychological models (Wetzels, Vandekerckhove, Tuerlinckx, and Wagenmakers, 2010), try to sequentially optimize the stimuli used in an experiment (Myung and Pitt, 2009), or model how participants learn to interact with their environment (Meder and Nelson, 2012), many problems require us to assess unknown functions that map inputs to outputs. Often, the shape of the underlying function is unknown, the function might be hard to evaluate analytically, or other requirements such as design costs might complicate the process of information acquisition. In these situations, Gaussian process regression can serve as a useful tool for performing inference both passively (for example, describing a given data set as best as possible, allowing one to also predict future data) as well as actively (for example, learning while choosing input points to produce the highest possible outputs, cf Williams and Rasmussen, 2006). Gaussian process regression is a non-parametric Bayesian approach (Gershman and Blei, 2012) towards regression problems. It can capture a wide variety of relations between inputs and outputs by utilizing a theoretically infinite number of parameters and letting the data determine the level of complexity through the means of Bayesian inference (Williams, 1998).

This tutorial will introduce Gaussian process regression as an approach towards describing, and actively learning and optimizing unknown functions. It is intended to be accessible to a general readership and focuses on practical examples and high-level explanations. It consists of six main parts: The first part will introduce the mathematical underpinnings of Gaussian process regression. The second part will show how different kernels can encode prior assumptions about the underlying function. Next, we will show how Gaussian processes can be used in problems of optimal experimental design, when the goal is pure exploration, i.e., to learn a function as well as possible. The fourth part will describe how Gaussian process-based Bayesian optimization (here defined as an *exploration-exploitation problem*) works. In the fifth part, we will talk about ways of utilizing Gaussian process exploration-exploitation methods in situations with additional requirements and show one example of “safe exploration”, where the goal is to avoid outputs below a certain threshold. We will conclude by summarizing current research that treats Gaussian process regression as a psychological model to assess human function learning.

As a tutorial like this can never be fully comprehensive, we have tried to provide detailed references and software pointers whenever possible.

## 2. Gaussian processes – distributions over functions

### 2.1. *Motivation*

Let *f* denote an (unknown) function which maps inputs *x* to outputs *y*: *f*: *X* ⟶ *Y*. Throughout the following examples, we will use Gaussian process regression to accomplish either one of three different goals:

By *modelling* a function *f* we mean mathematically representing the relation between inputs and outputs. An accurate model of *f* allows us to predict the output for many possible input values. In practice, this means collecting observations of both inputs and outputs and on the basis of this generating accurate predictions for newly observed points. As an example of this, we will use Gaussian process regression to model mouse trajectories in a categorization experiment. Additionally, we will use compositional Gaussian process regression to decompose temporal dependencies in participants’ reaction times into interesting patterns.

By *exploring* a function we mean to actively choose the input points for which to observe the outputs in order to accurately model the function. In pure exploration problems, the only objective is to explore the underlying function well in order to learn about it as quickly and accurately as possible. This set-up is closely related to optimal experimental design scenarios as it equates to adaptively selecting the input points based on what is already known about the function and where knowledge can be improved. In a simple simulation experiment, we will show how exploration based on Gaussian process regression can recover underlying response functions faster than other commonly used techniques.

In *exploration-exploitation* problems, the outcomes of chosen inputs are accrued over time. The objective is to find inputs that produce the highest outputs in order to maximise the total reward accrued within a particular period of time. Exploration solely serves the purpose of doing so most effectively. This set-up is closely related to optimization problems as the goal is to find the maximum of the function as efficiently as possible. It is called *exploration-exploitation* as scenarios where the output of the underlying function has to be optimized require us to both sample uncertain areas in order to gain more knowledge about the function (exploration) as well as sampling input points that are likely to generate high outputs given the current knowledge of the function (exploitation). As an example, we will show how Gaussian process-based exploration-exploitation quickly finds highly rated items in a movie recommendation application. Moreover, we will show how this method can be adapted to additional requirements such as avoiding outputs below a given threshold.

Both *exploration* and *exploration-exploitation* tasks require choosing useful inputs. Doing so requires two ingredients:

1. A model used to learn about the function *f*.
2. A method to select inputs based on the current knowledge of *f*.

As a valid model of the underlying function *f* is crucial for all three goals of modelling, exploration, and exploitation, we will first focus on Gaussian processes as a powerful and expressive method to model unknown functions. We will focus on applying this tool to exploration-exploitation scenarios afterwards. Table 1 provides an overview of the different Gaussian process methods (and their example applications) introduced in this tutorial.

**Table 1:** Overview of different Gaussian process methods (including their example applications) introduced in this tutorial.

| Method | Purpose | Approach | Example |
| --- | --- | --- | --- |
| Modelling | Simple regression | passive | Mouse trajectories |
| Compositional<br>modelling | Find patterns within<br>data | passive | Response time patterns |
| Exploration | Learn function as<br>quickly as possible | active | Learn simulated<br>functions |
| Exploration-exploitation | Optimize function | active | Movie recommendation |
| Safe<br>exploration | Optimize function<br>while staying above a<br>threshold | active | Cautious stimulus<br>optimization |

**Table 2:** Observations for the regression example. Inputs *x*_*t*_ and corresponding outputs *y*_*t*_ observed at 6 different times *t* = 1,…, 6.

| $t$ | $x_t$ | $y_t$ |
| --- | --- | --- |
| 1 | 0.9 | 0.1 |
| 2 | 3.8 | 1.2 |
| 3 | 5.2 | 2.1 |
| 4 | 6.1 | 1.1 |
| 5 | 7.5 | 1.5 |
| 6 | 9.6 | 1.2 |

### 2.2. *Modelling functions: the weight space view*

Let us start by considering a standard approach to model functions: linear regression (here approached from a Bayesian viewpoint). Imagine we have collected the observations shown in Table 2 and that we want to predict the value of *y* for a new input point *x*_⋆_ = 3. In linear regression, we assume that the outputs are a linear function of the inputs with additional noise:

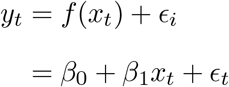

where the noise term *∊*_*t*_ follows a normal distribution

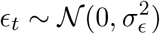

with mean 0 and variance of 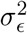. As this will be useful later, we can also write this in matrix algebra as

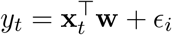

defining the vectors

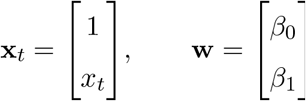

To predict the output for *x*_⋆_, we need to estimate the weights from the previous observations

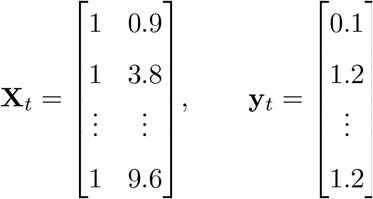

Adopting a Bayesian framework, we do so through the posterior distribution over the weights. If we use a Gaussian prior over the weights 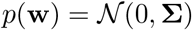 and the Gaussian likelihood 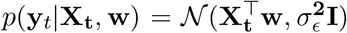, then this posterior distribution is

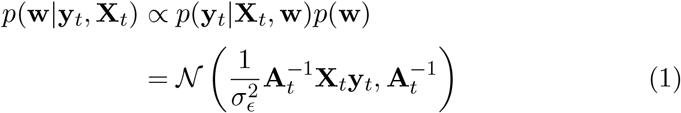

where 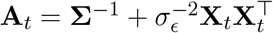 (see also Williams, 1998).

As inference is performed over the weights (i.e., we try to find the best estimate for the *β*-weights given the data), this is also sometimes referred to as “the weight space view of regression”. To predict the output *y*_⋆_ at a new test point **x**_⋆_, we can average out the error term and focus on the expected value which is provided by the function *f*, predicting *f*_⋆_ = *y*_⋆_ – ∊_⋆_ = *f* (x_⋆_). In the predictive distribution of *f*_⋆_, we average out our uncertainty regarding the weights

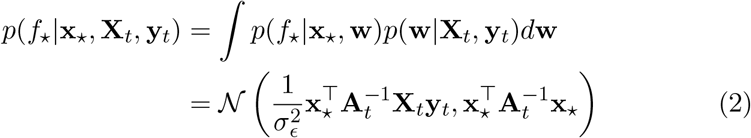

You can also imagine generating this posterior predictive distribution over *f*_⋆_ by first sampling weights from the posterior distribution over weights (see Equation 1), and then using these sampled weights to generate predictions for the new input points.

A good point prediction of *y*_⋆_ is the mean of this predictive distribution. Comparing the mean in (2) to the mean in (1), we see that we can simply multiply the posterior mean of **w** with the new input **x**_⋆_, resulting in the prediction 0.56 + 3 × 0.12 = 0.92.

While linear regression is often chosen to model functions, it assumes the function has indeed a linear shape. However, only few relations in the real world are truly linear, and we need a way to model non-linear dependencies as well. One possible adjustment is to use a mapping of the inputs **x** onto a “feature space”, i.e. by transforming the inputs with a non-linear function *Φ*(**x**), resulting in an *n*-dimensional vector of numerical features representing the transformed input. After transformation, we can again perform linear Bayesian regression, but now on the transformed input. A common mapping is to use polynomials, resulting in polynomial regression. Take cubic regression as an example, which assumes a function *f*(*x*) = *β*_0_ + *β*_1_*x* + *β*_2_*x*^2^ + *β*_3_*x*^3^. Deriving the posterior for this model is similar to the linear regression described before, only that the input matrix **X**_*t*_ is replaced by the mapping:

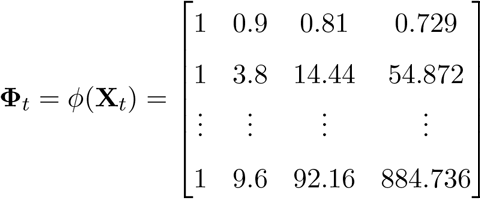

In our example –and again using the posterior mean of the weights– this would result in the prediction *f*_⋆_ = −0.67 + 0.98 × 3 — 0.13 × 3^2^ + 0.01 × 3^3^ = 1.37.

**Figure 1:**
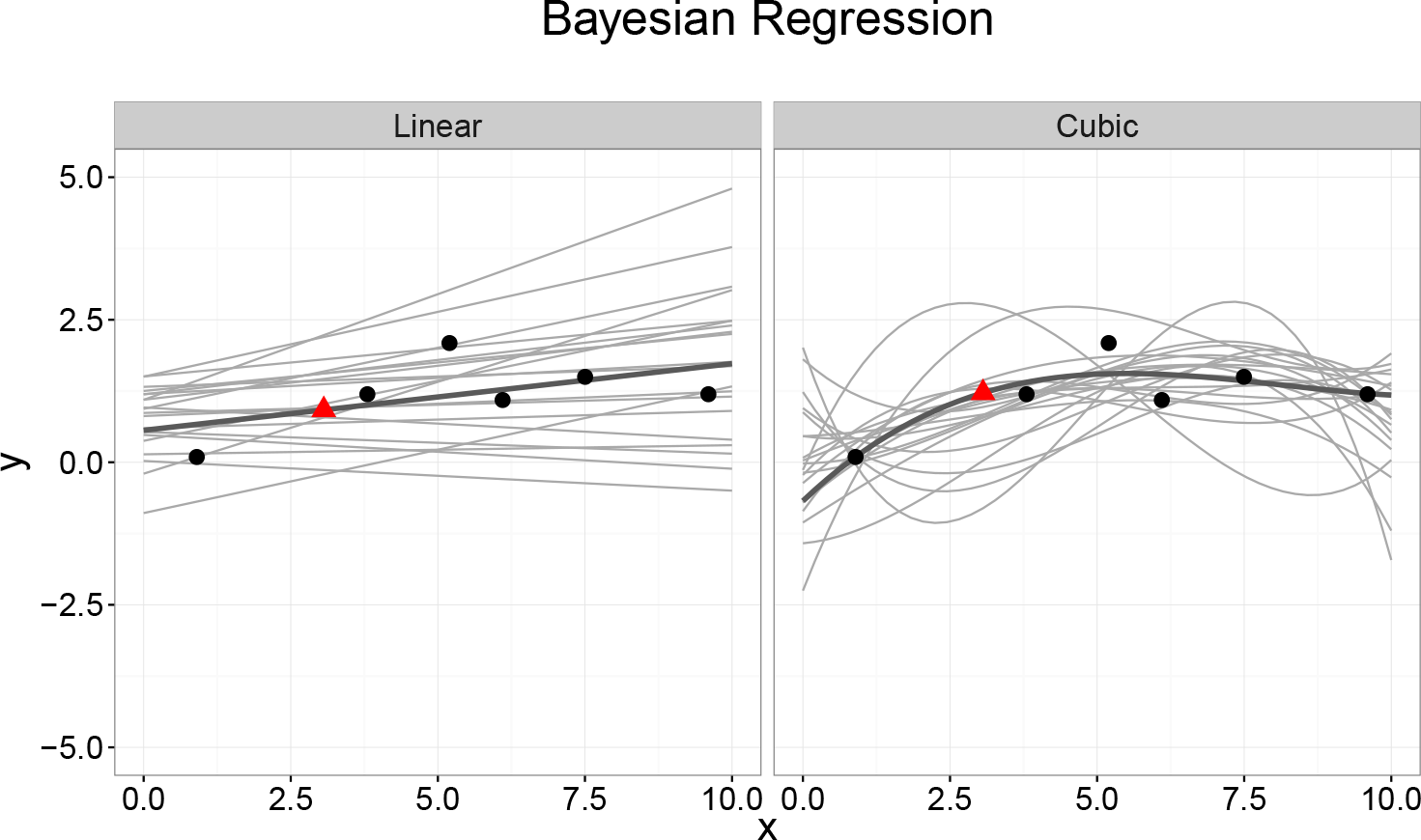
Example of performing Bayesian linear and cubic regression. Grey lines indicate predictions for different sampled posterior weights. Black dots mark empirical observations. Dark grey lines mark the current mean posterior predictions. The red triangle shows the prediction for a new data point *x*_⋆_ = 3.

Mapping input variables into a feature space offers considerably more flexibility and allows one to model functions of any shape. However, this flexibility is also a drawback. There are infinitely many mappings possible and we have to choose one either a priori or by model comparison within a set of possible mappings. Especially if the problem is to explore and exploit a completely unknown function, this approach will not be beneficial as there is little guidance to which mapping we should try. Gaussian process regression, to which we turn next, offers a principled solution to this problem in which mappings are chosen implicitly, effectively letting “the data decide” on the complexity of the function^1^

## 2.3. *Modelling functions: the function space view*

In the weight space view of the previous section, we focused on distributions over weights. As each set of weights implies a particular function, a distribution over weights implies a distribution over functions. In Gaussian process regression, we focus directly on such distributions over functions.

A Gaussian process defines a distribution over functions such that, if we pick any two or more points in a function (i.e., different input-output pairs), observations of the outputs at these points follow a joint (multivariate) Gaussian distribution. More formally, a Gaussian process is defined as a collection of random variables, any finite number of which have a joint (multivariate) Gaussian distribution.

In Gaussian process regression, we assume the output *y* of a function *f* at input x can be written as

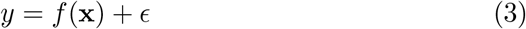

with 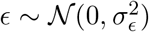. Note that this is similar to the assumption made in linear regression, in that we assume an observation consists of an independent “signal” term *f*(**x**) and “noise” term ∊. In Gaussian process regression, however, we assume that the signal term is also a random variable which follows a particular distribution. This distribution is subjective in the sense that the distribution reflects our uncertainty regarding the function. The uncertainty regarding *f* can be reduced by observing the output of the function at different input points. The noise term ∊ reflects the inherent randomness in the observations, which is always present no matter how many observations we make. In Gaussian process regression, we assume the function *f*(**x**) is distributed as a Gaussian process:

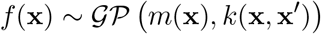

A Gaussian process 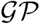 is a distribution over functions and is defined by a *mean* and a *covariance* function. The mean function *m*(**x**) reflects the expected function value at input **x**:

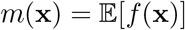

i.e. the average of all functions in the distribution evaluated at input **x**. The prior mean function is often set to *m*(**x**) = 0 in order to avoid expensive posterior computations and only do inference via the covariance function. Empirically, setting the prior to 0 is often achieved by subtracting the (prior) mean from all observations. The covariance function *k*(**x**, **x′**) models the dependence between the function values at different input points **x** and **x′**:

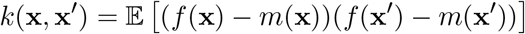

The function *k* is commonly called the *kernel* of the Gaussian process (Jäkel, Schölkopf, and Wichmann, 2007). The choice of an appropriate kernel is based on assumptions such as smoothness and likely patterns to be expected in the data. A sensible assumption is usually that the correlation between two points decays with the distance between the points. This means that closer points are expected to behave more similarly than points which are further away from each other. One very popular choice of a kernel fulfilling this assumption is the radial basis function kernel, which is defined as

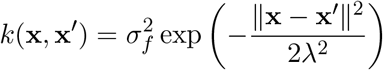

The radial basis function provides an expressive kernel to model smooth and stationary functions. The two hyper-parameters λ (called the length-scale) and 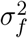 (the signal variance) can be varied to increase or reduce the a priori correlation between points and consequentially the variability of the resulting function.

Once a mean function and kernel are chosen, we can use the Gaussian process to draw a priori function values, as well as posterior function values conditional upon previous observations.

### 2.3.1. *Sampling functions from a GP*

Although Gaussian processes are continuous, sampling a function from a Gaussian process is generally done by computing the function values of a selected set of input points. Theoretically, a function can be represented as a vector of infinite size; however, as we only have to make predictions for finitely many points in practice, we can draw outputs for these points by using a multivariate normal distribution with a covariance matrix generated by the kernel. Let **X**_⋆_ be a matrix with on each row a new input point 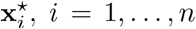. To sample a function, we first compute the covariances between all inputs in **X**_⋆_ and collect these in an *n* × *n* matrix:

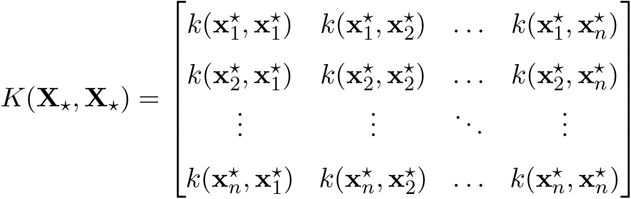

Choosing the usual prior mean function *m*(**x**) = 0 to simplify the matrix algebra shown in Equation 4, we can then sample values of *f* at inputs **X**_⋆_ from the 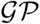 by sampling from a multivariate normal distribution

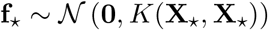

where we use the notation 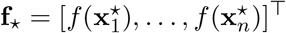. Note that **f**_⋆_ is a sample of the function values. To sample observations **y**_⋆_, we would have to add an additional and independent sample of the noise term ∊.

### 2.3.2. *Posterior predictions from a GP*

Suppose we have collected observations 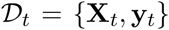 and we want to make predictions for new inputs **X**_⋆_ by drawing **f**_⋆_ from the posterior distribution 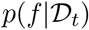. By definition, previous observations **y**_*t*_ and function values **f**_⋆_ follow a joint (multivariate) normal distribution. This distribution can be written as

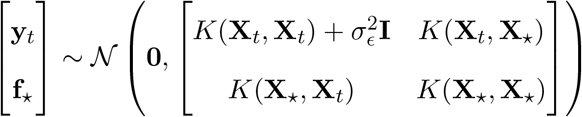

where *K*(**X**_*t*_, **X**_*t*_) is the covariance matrix between all observed points so far, *K*(**X**_⋆_, **X**_⋆_) is the covariance matrix between the newly introduced points as described earlier, *K*(**X**_⋆_, **X**_*t*_) is the covariance matrix between the new input points and the already observed points and *K*(**X**_*t*_, **X**_⋆_) is the covariance matrix between the observed points and the new input points. Moreover, I is an identity matrix (with 1’s on the diagonal, and 0’s elsewhere) and 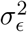 is the assumed noise level of observations (i.e. the variance of ∊). Using standard results (see for example Rasmussen and Nickisch, 2010), the conditional distribution *p*(**f**_⋆_|**X**_*t*_, **y**_*t*_, **X**_⋆_) is then a multivariate normal distribution with mean

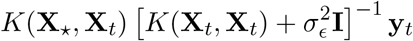

and covariance matrix

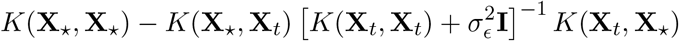

Note that this posterior is also a GP with mean function\

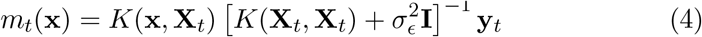

and kernel

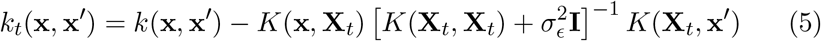

This means that calculating the posterior mean and covariance of a GP involves first calculating the 4 different covariance matrices above and then combining them according to Equations 4-5. In order to aid the understanding of the matrix algebra involved in these calculations, the different matrices are represented visually in Figure 2.

**Figure 2:**
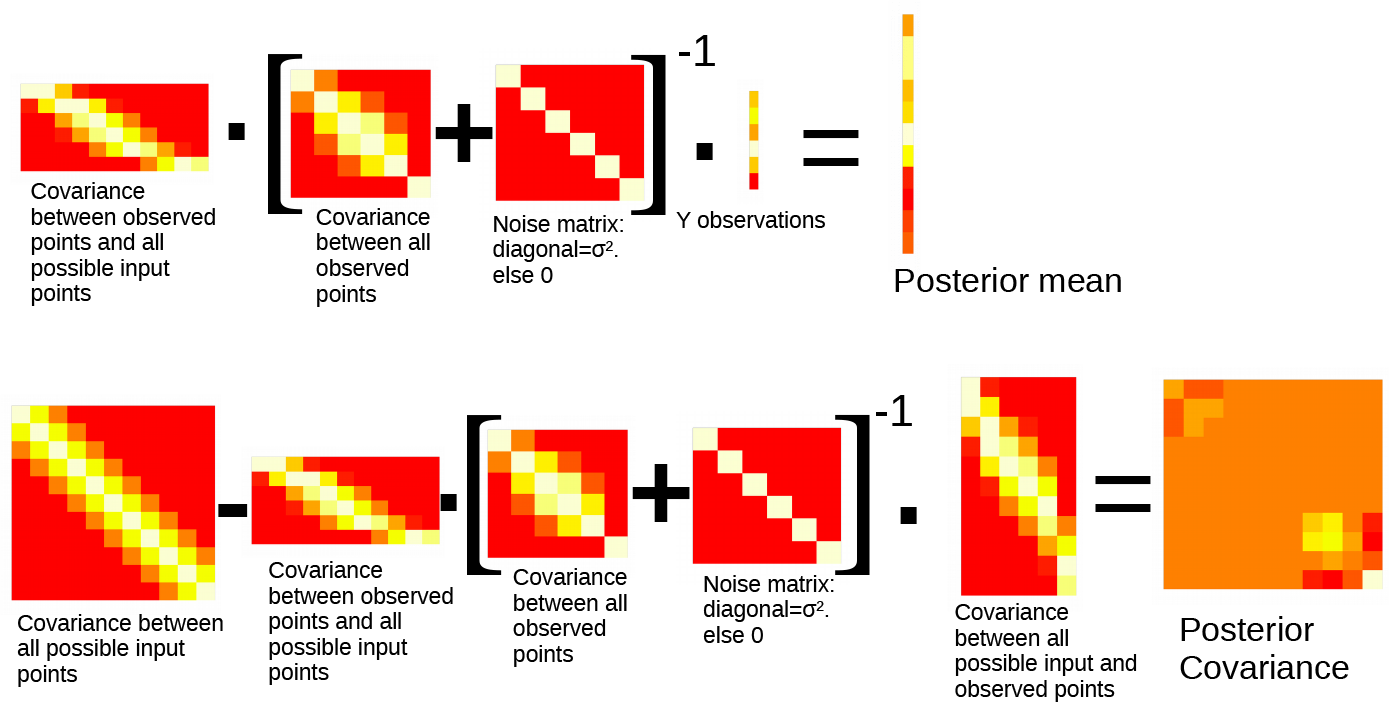
Visual representation of calculating the GP posterior mean and covariance given the example points from Table 2. Lighter colours indicate higher values. For the posterior mean, the covariance between all observed points is multiplied by the inverse of the sum of the covariance of the observed points and the noise matrix, as well as by the observations of the dependent variable. For the posterior covariance, the overall covariance between all possible input points is calculated and afterwards the product of the covariance between the observed points and all possible input points, the inverse of the sum between the covariance of the observed points and the noise matrix, as well as the covariance between all possible input points and the observed points, is subtracted.

To predict **f**_⋆_, we can simply use the mean function in (4), or sample functions from the GP with this mean function and the kernel in (5), as described in the previous section.

Figure 3 shows an example of samples from a GP prior with a radial basis function kernel (with λ = 0.5), and samples from the posterior mean functions after the data in Table 2 has been observed.

### 2.3.3. *Switching back to the weight view*

We can rewrite the mean function in (4) as

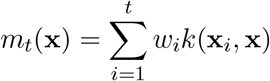

where each **x**_*i*_ is a previously observed input value in **X**_*t*_ and the weights are collected in the vector 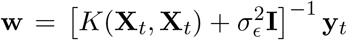. This equation shows that Gaussian process regression is equivalent to a linear regression model using basis functions *k* to project the inputs into a feature space. To make new predictions, every output *y*_*t*_ is weighted by how similar its associated input **x**_*t*_ is to the to-be-predicted point **x** by a similarity measure induced by the kernel. This results in a simple weighted sum to make predictions for new points^2^. Therefore, a conceptually infinite parameter space boils down to a finite sum when making predictions ^3^. This sum only depends on the chosen kernel *k* and the data 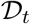 observed thus far (Kac and Siegert, 1947). This is why Gaussian process regression is referred to as a non-parametric technique. It is not the case that this regression approach has no parameters; actually, it has theoretically as many parameters *w* as there are observations. However, making predictions involves only a finite sum over all past observations. Details for generating a prediction for *x*_⋆_ = 3 given a radial basis function kernel with length scale λ = 1, observation variance 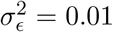, and signal variance 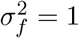 are provided in Table 4.

**Figure 3:**
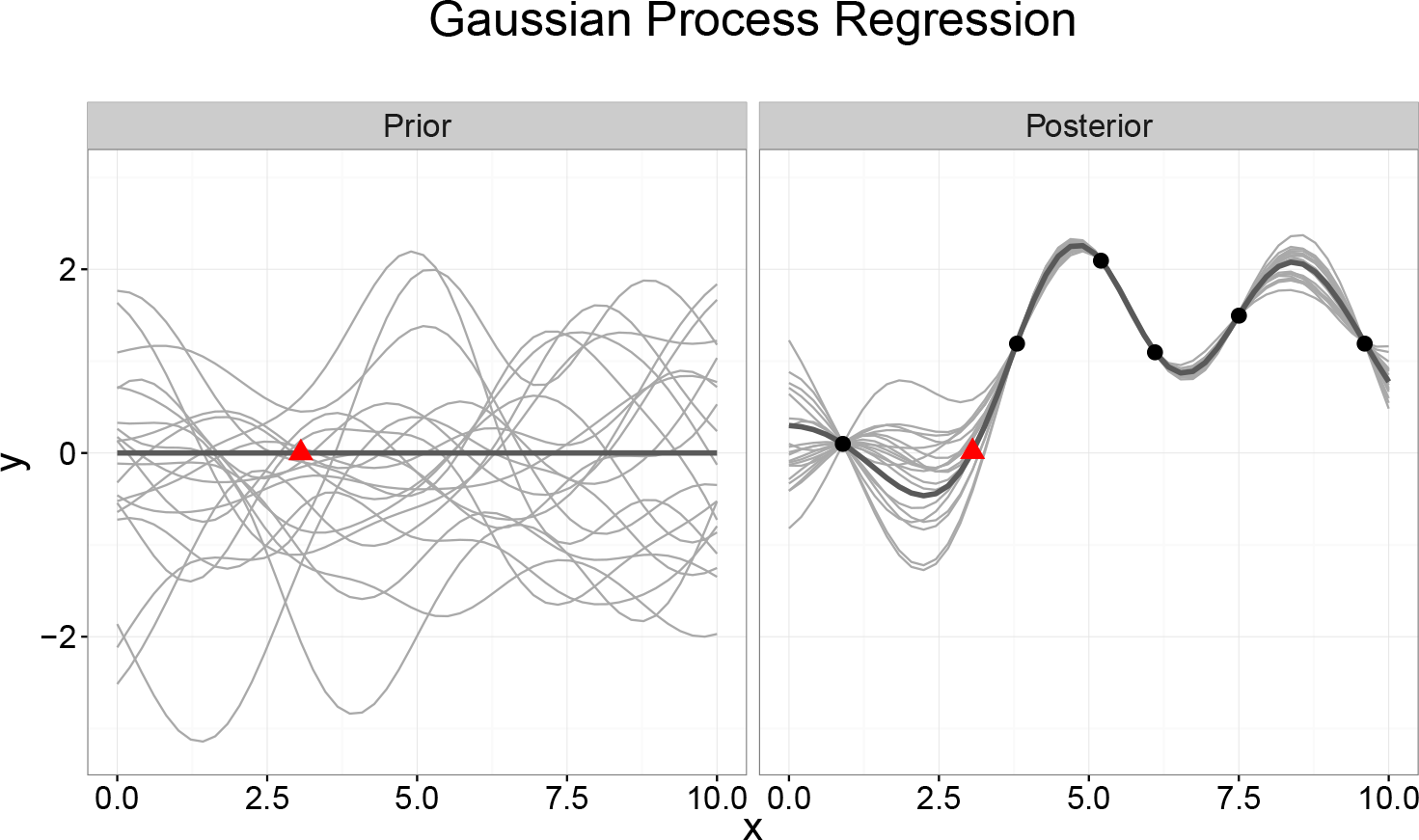
Samples from a Gaussian process prior and posterior. Grey lines indicate samples from the GP. Black dots mark empirical observations. The dark grey line marks the current mean of the GP. The red triangle shows the prediction for the new input point.

**Table 3.** Example of generating a prediction using a Gaussian process with a radial basis function kernel 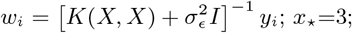

| $t$ | $x_t$ | $y_t$ | $w_t$ | $k(x_t, x_\star)$ | $w_t k(x_t, x_\star)$ |
| --- | --- | --- | --- | --- | --- |
| 1 | 0.9 | 0.1 | 0.51 | 0.38 | 0.19 |
| 2 | 3.8 | 1.2 | -3.88 | 0.87 | -3.37 |
| 3 | 5.2 | 2.1 | 13.3 | 0.34 | 4.53 |
| 4 | 6.1 | 1.1 | -12.55 | 0.12 | -1.48 |
| 5 | 7.5 | 1.5 | 5.83 | 0.01 | 0.06 |
| 6 | 9.6 | 1.2 | -0.34 | 0.00 | 0.00 |
| | | | | $\sum_{t=1}^6 w_t k(x_t, x_\star)$: | -0.06 |

### 2.4. *Optimizing hyper-parameters*

The kernel usually contains hyper-parameters such as the length-scale, signal variance, and noise variance, which are unknown and need to be inferred from the data. As the posterior distribution over the hyper-parameters is non-trivial to obtain, full Bayesian inference of the hyper-parameters is not frequently used in practice. Instead, common practice is to obtain point estimates of the hyper-parameters by maximising the marginal (log) likelihood. This is similar to parameter estimation by maximum likelihood and is also referred to as type-II maximum likelihood (ML-II, cf Williams and Rasmussen, 2006). Given the data 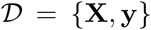 and hyper-parameters ***θ*** (e.g.,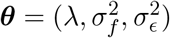), the log marginal likelihood is

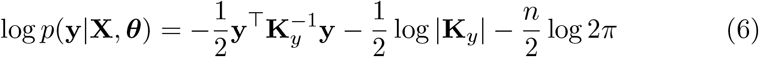

where 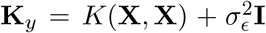 is the covariance matrix of the noisy output values *y*. The marginal log likelihood can be viewed as a penalized fit measure, where the term 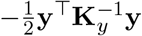 measures the data fit –that is how well the current kernel parametrization explains the dependent variable-and 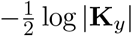 is a complexity penalization term. The final term 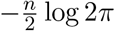 is a normalization constant. The marginal likelihood is normally maximized through a gradient-ascent based optimization tool such as implemented in Carl Rasmussen’s MATLAB function `minimize.m`^4^. These routines make use of the partial derivatives of (6) with respect to ***θ***:

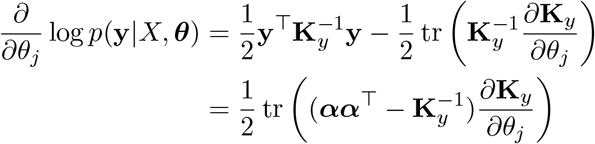

with 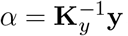.

There are recent efforts to make hyper-parameter estimation fully Bayesian, for example by using `Stan` (Flaxman, Gelman, Neill, Smola, Vehtari, and Wilson, 2015), which are promising to result in more robust estimates by additionally providing uncertainty estimates for the obtained parameters.

## 3. Example: Modelling mouse trajectories

As an example of modelling functions, we consider mouse trajectory data from Kieslich and Henniger (2017). Participants performed a task in which they had to classify animals (for example, a lion or a falcon) into different categories (for example, a mammal or a bird) by using a computer mouse to move a cursor from a start position (on the left of the screen) to the correct category (on the right of the screen). Kieslich and Henniger tracked the location of participants’ cursors at different time points, and these discretized points can be summarized by functions describing movement trajectories over the screen. Studying mouse trajectory data can reveal additional realtime information about psychological processes such as categorization and perception (see Freeman and Ambady, 2010).

Gaussian process regression has been successfully applied to such scenarios, where it is useful as the priors over different functions can also be modelled hierarchically, thereby assessing whether participants move the mouse differently for typical (e.g., “monkey-mammal”) or atypical (e.g., “penguin-bird”) category members, as described in more detail by Cox, Kachergis, and Shiffrin (2012). Here, we simply want to test if Gaussian process regression can be used as an appropriate smoothing technique for such data. Smoothing mouse trajectory data is especially important if one wants to make claims about the underlying shapes of group-level trajectories, for example whether or not trajectories look different for typical than for atypical exemplars. Additionally, smoothing mouse trajectories by using Gaussian process regression comes with the additional benefit that possible posterior trajectories can be samples as the GP provides not only a descriptive but also a generative model of the data.

We take participants’ raw trajectory data (their x-y-coordinates) over time and assess how well Gaussian process regression is able to predict left-out trajectory points. More specifically, we use participants *x* coordinates as input, and the *y* coordinates as output; for every trajectory, we randomly sample 80% of the points and use them as a training set, and then predict the left-out 20% trajectory points. In order to make meaningful claims about GP’s usefulness, we compare its performance to two other smoothing techniques. First, a polynomial regression with up to 5 degrees, where the order is chosen by Akaike’s “An Information Criterion” (Akaike, 1974; Lee, 2004). Secondly, a cubic smoothing spline with the degrees of freedom determined by cross validation within the training set (Durrleman and Simon, 1989).

The left part of Figure 4 shows the mean square error over 1000 runs including the attached standard error. We can see that Gaussian process regression produces a lower out-of-sample prediction error than either the polynomial regression or the spline smoothing, thereby demonstrating that it is a useful tool for mouse trajectory modelling. The right part of Figure 4 shows an example of smooth lines generated by Gaussian process regression.

## 4. Encoding prior assumptions via the kernel

So far we have only focused on the radial basis function kernel to perform Gaussian process regression. However, other kernels are possible and flexibility in choosing the kernel is one of the benefits of Gaussian process regression. The kernel function *k* directly encodes prior assumptions about the underlying function such as its smoothness and periodicity. Additionally, more complex kernels can be created by combining simpler kernels through operations such as addition or multiplication.

### 4.1. *Encoding smoothness*

The radial basis function kernel is a special case of a general class of kernel functions called the Matérn kernel. The Matérn covariance between two points with distance τ = |x – x′| is

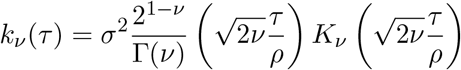

where Γ is the gamma function, *K*_*v*_ is the modified Bessel function of the second kind, and *ρ* and *ν* are non-negative covariance parameters. A 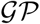 with a Matérn covariance function has sample paths that are *ν* – 1 times differentiable. When *ν* = *p* + 0.5, the Matérn kernel can be written as a product of an exponential and a polynomial of order *p*.

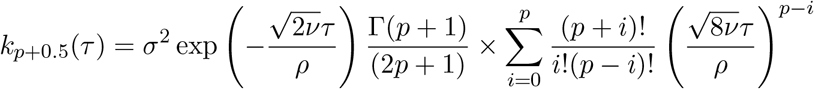

**Figure 4:**
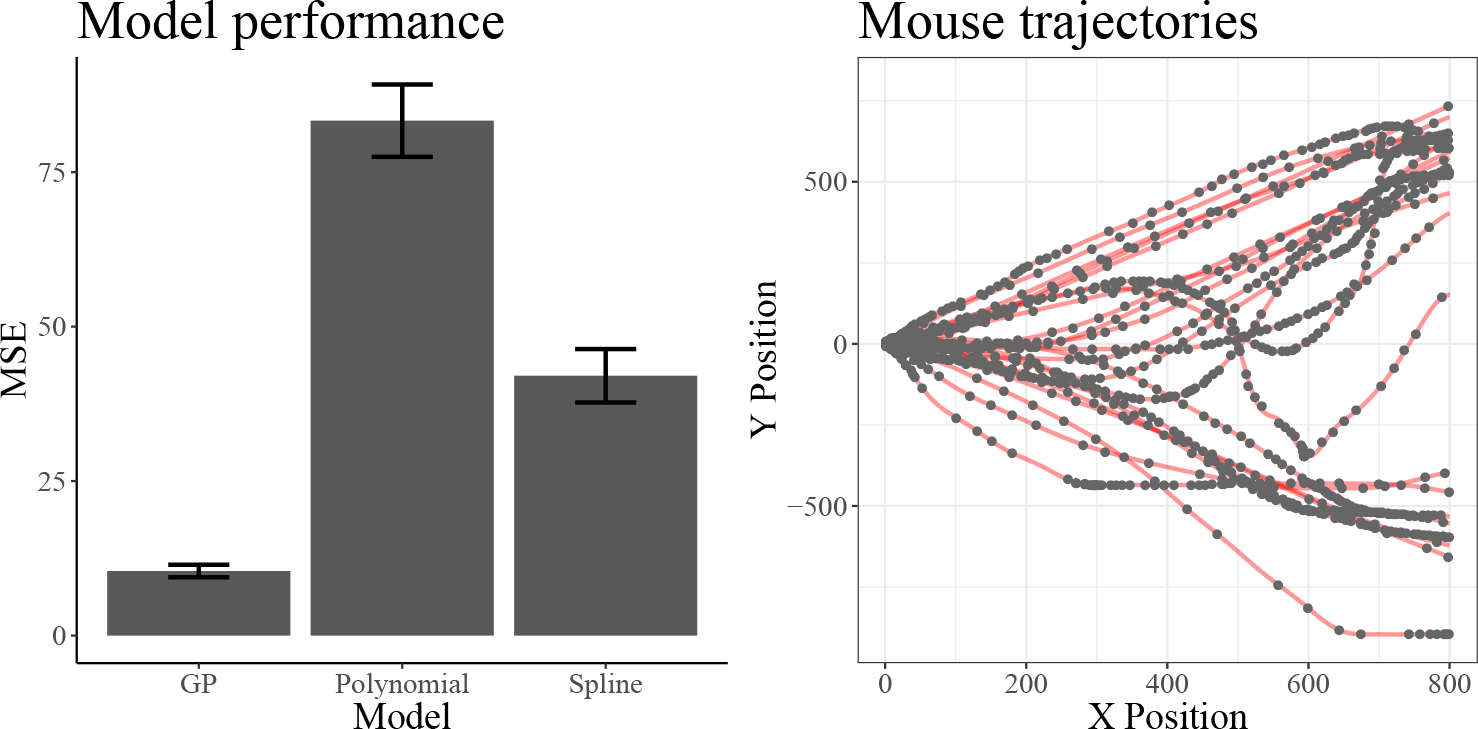
Modeling mouse trajectories. **Left:** Performance of Gaussian Process mouse trajectory smoothing as compared to Splines and polynomial regression. Error bars represent the standard error of the mean. **Right:** Smoothing lines produced by Gaussian Process regression.

Here, *p* directly determines how quickly the covariance between two points thins out in dependency of the distance between the two points. If *p* = 0, then this leads to the Ornstein-Uhlenbeck process kernel

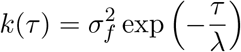

which encodes the prior assumption that the function is extremely unsmooth (rough) and that observations do not provide a lot of information about points that are anything but very close to the points we have observed so far. In the limit as *p* ⟶ ∞, the Matérn kernel becomes a radial basis function kernel. This kernel expects very smooth functions for which observing one point provides considerably more information than if we assume very rough underlying functions. Figure 5 shows prior and posterior samples for both the Ornstein-Uhlenbeck process and the radial basis function kernel. Notice how the prior samples are a lot more “rugged” for the former and very smooth for the later. We can also see how encoding different prior smoothness assumptions leads to different posterior samples after having observed the same set of points (the points we used before). In particular, expecting very rough functions a priori leads to posteriors that do not generalize far beyond the encountered observations, whereas expecting very smooth functions leads to posterior samples that generalize more broadly beyond the encountered points.

In most real world applications, practitioners choose the radial basis function kernel and then optimize its length-scale in order to account for potential mismatches between prior smoothness assumptions and the observed data. The main reason for this is that the radial basis function kernel is easy to specify and also computationally convenient as one only has to evaluate an exponentiated distance instead of a product between a polynomial and an exponent as is the case for the Matérn kernel. Within exploration-exploitation scenarios, another frequent choice is to use a Matérn kernel with *p* = 5 as an intermediate solution to encode the expectation of “smooth but not too smooth” functions. However, instead of relying on such default choices, it will be usually better to choose the level of smoothness by considering the expected properties of the underlying function, in order to avoid mismatched priors (Schulz, Speekenbrink, Hernández-Lobato, Ghahramani, and Gershman, 2016c). For example, whereas mouse trajectories are normally smooth and therefore might lend themselves well to using a radial basis function kernel, other processes such as eye movements might be less smooth and therefore modelled more precisely with an Ornstein-Uhlenbeck kernel (Engbert and Kliegl, 2004).

**Figure 5:**
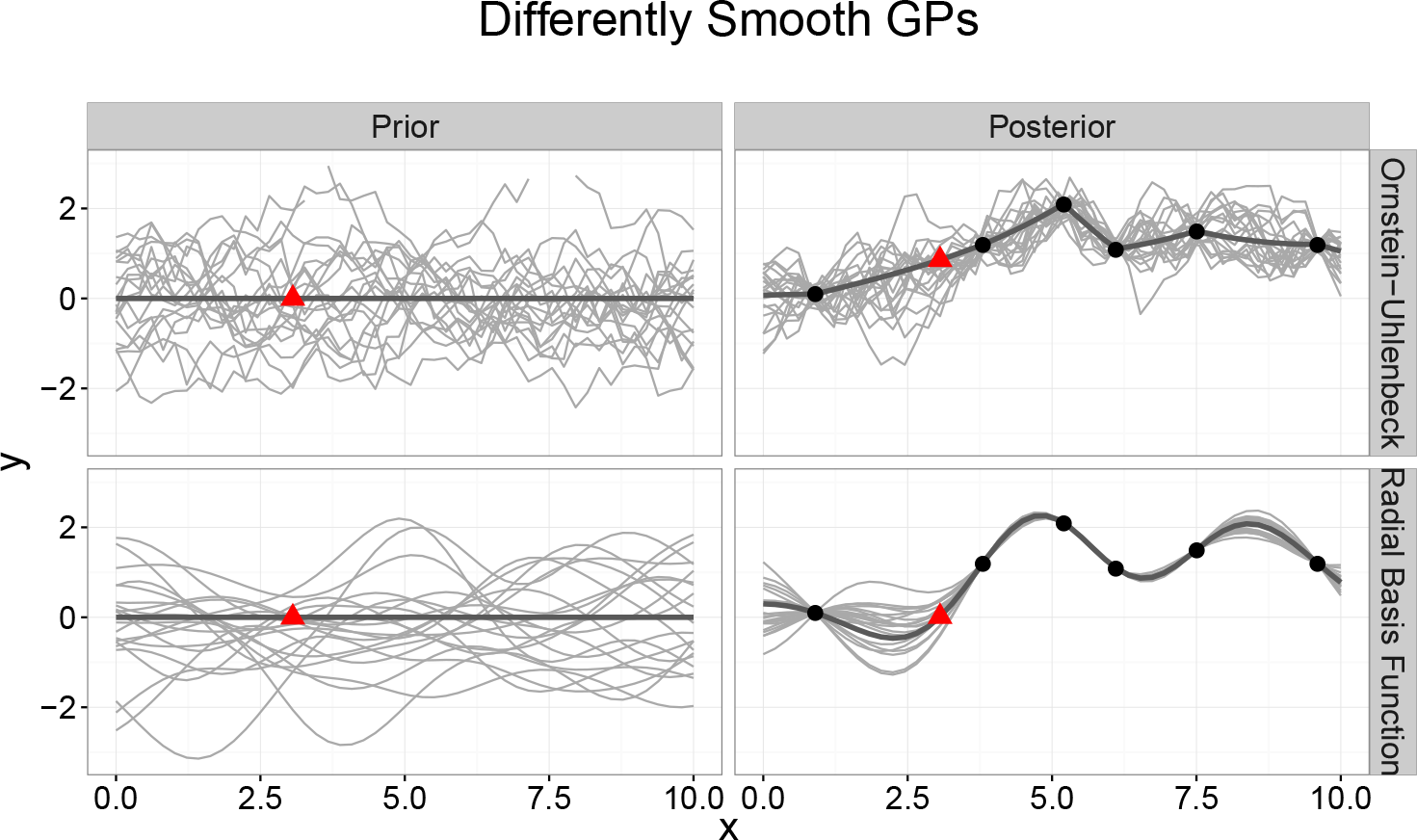
Samples from differently smooth Gaussian process priors and posteriors after having observed the same set of points. Grey lines indicate samples from the GP. Black dots mark empirical observations. The dark grey line marks the current mean of the GP. The red triangle shows the prediction for the new data point.

### 4.2. *Composing kernels*

Another advantage of Gaussian process regression is that different kernels can be combined, thereby creating a rich set of interpretable and reusable building blocks (Duvenaud, Lloyd, Grosse, Tenenbaum, and Ghahramani, 2013). For example, adding two kernels together models the data as a superposition of independent functions. Multiplying a kernel with a radial basis function kernel, locally smooths the predictions of the first kernel.

Take the data set of atmospheric concentration of carbon dioxide over a forty year horizon as shown in Figure 6. We can immediately see a pattern within this data, which is that the CO2-concentration seems to increase over the years, that there seems to be some periodicity by which at some times within each year the CO2 emission is higher, and that this period may not be perfectly replicated every year. Using a Gaussian process regression framework, we can combine different kernels as building blocks in the attempt to explain these patterns. Figure 6 shows posterior mean predictions for different kernel combinations.

**Figure 6:**
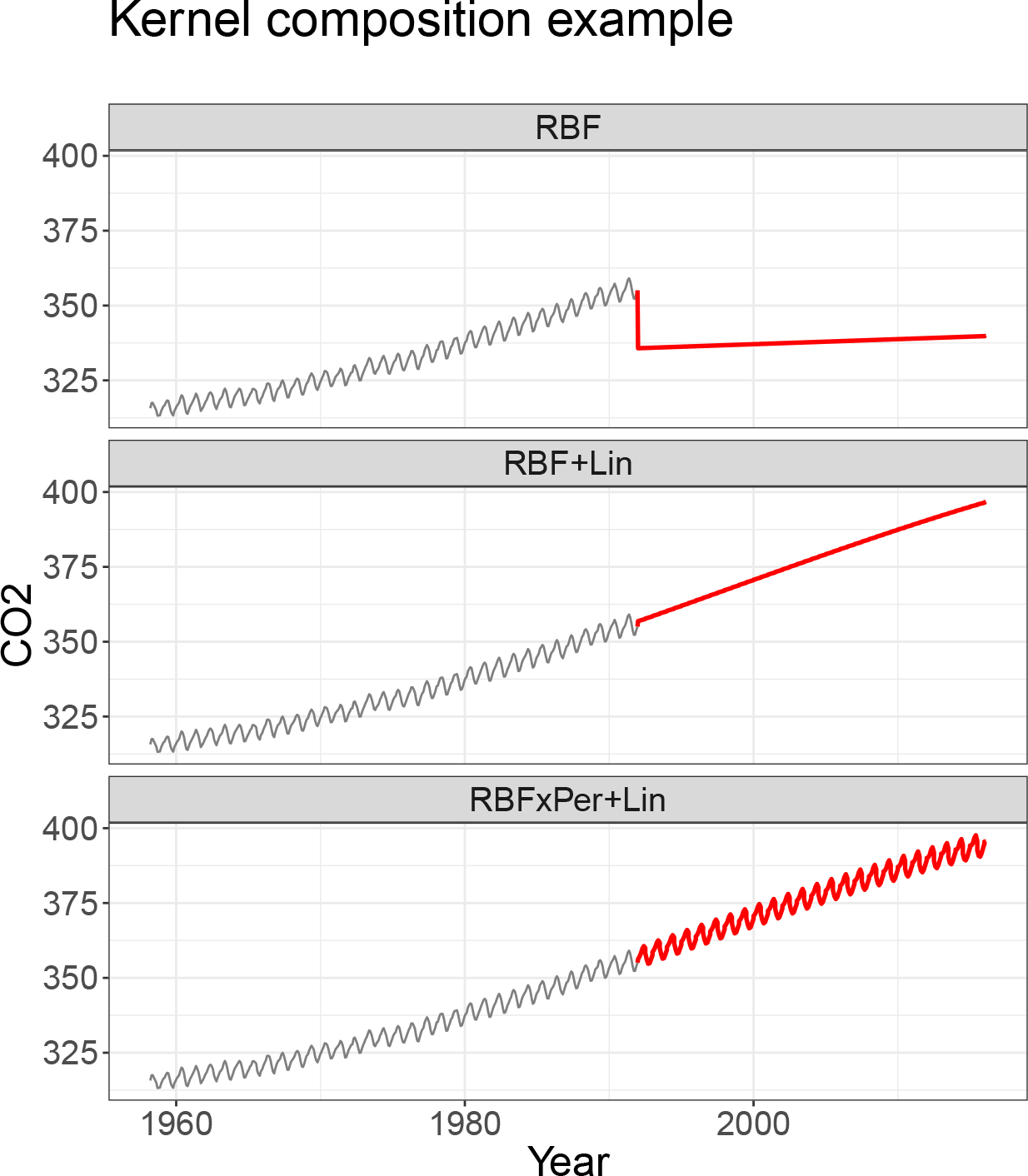
Example of composing kernels by combining simpler kernels in order to explain a complex function. Data were mean-centred before fitting the Gaussian process and predictions were transformed back afterwards. Grey lines show observed CO2 emissions. Red lines show posterior predictions of Gaussian process regressions with different kernels: RBF is a radial basis function kernel, RBF+Lin is a kernel composed by adding a RBF and a linear kernel, RBF×Per + Lin is a kernel composed by multiplying a radial basis and periodic kernel and adding a linear kernel.

The first one shows a radial basis function alone, the second a sum of a radial basis function kernel and a linear kernel, *k*(*x*,*x*′) = (*x* – *c*)(*x*′ – *c*), and the third one the sum between a linear kernel and the product between a radial basis function kernel and a periodic kernel, 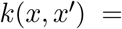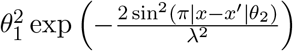. As the radial basis function kernel tends to reverse back to the mean over time, it does not do a good job capturing the linear trend of the data. Therefore, adding a linear kernel to the radial basis function kernel already seems to improve predictions. Finally, multiplying the radial basis function kernel with a periodic kernel to create a locally smoothed periodic kernel, which is then combined with an increasing trend by adding a linear kernel seems to predict the data best. This shows that the kernel can also be used to encode structural assumptions about the underlying function more explicitly, especially when one wants to cover more complex patterns than just interpolating smooth functions. Lloyd, Duvenaud, Grosse, Tenenbaum, and Ghahramani (2014) show how compositional Gaussian process regression can be used to create an “automatic statistician” which generates a full descriptive report when provided with a time series.

### 4.3. *Example: Temporal dependencies in response time analysis*

Compositional Gaussian process regression is most useful if the underlying function is supposed to show some inherent structure. One application for which structural patterns have been discussed in the literature is the analysis of long distance dependencies of response time patterns (Wagenmakers, Farrell, and Ratcliff, 2004; Van Zandt and Townsend, 2014). In particular, previous investigation suggest that response times over multiple trials are dependent based on an auto-regressive term (i.e., previous response times can predict the following) and a moving average term (i.e., the average response time can shift over trials). Here, we use compositional Gaussian Process regression in order to see what kind of patterns it can extract from participants long distance response time trials. For this, we analyse 4 participants of Wagenmakers et al. (2004) original study investigating long distance dependencies. We do not think that this analysis can supplant the more detailed approaches described in the literature, but nonetheless think it is interesting to probe such data sets for compositional patterns.

The results of a compositional Gaussian process regression modelling response times over 500 trials are shown for each participant individually in Figure 7.

**Figure 7:**
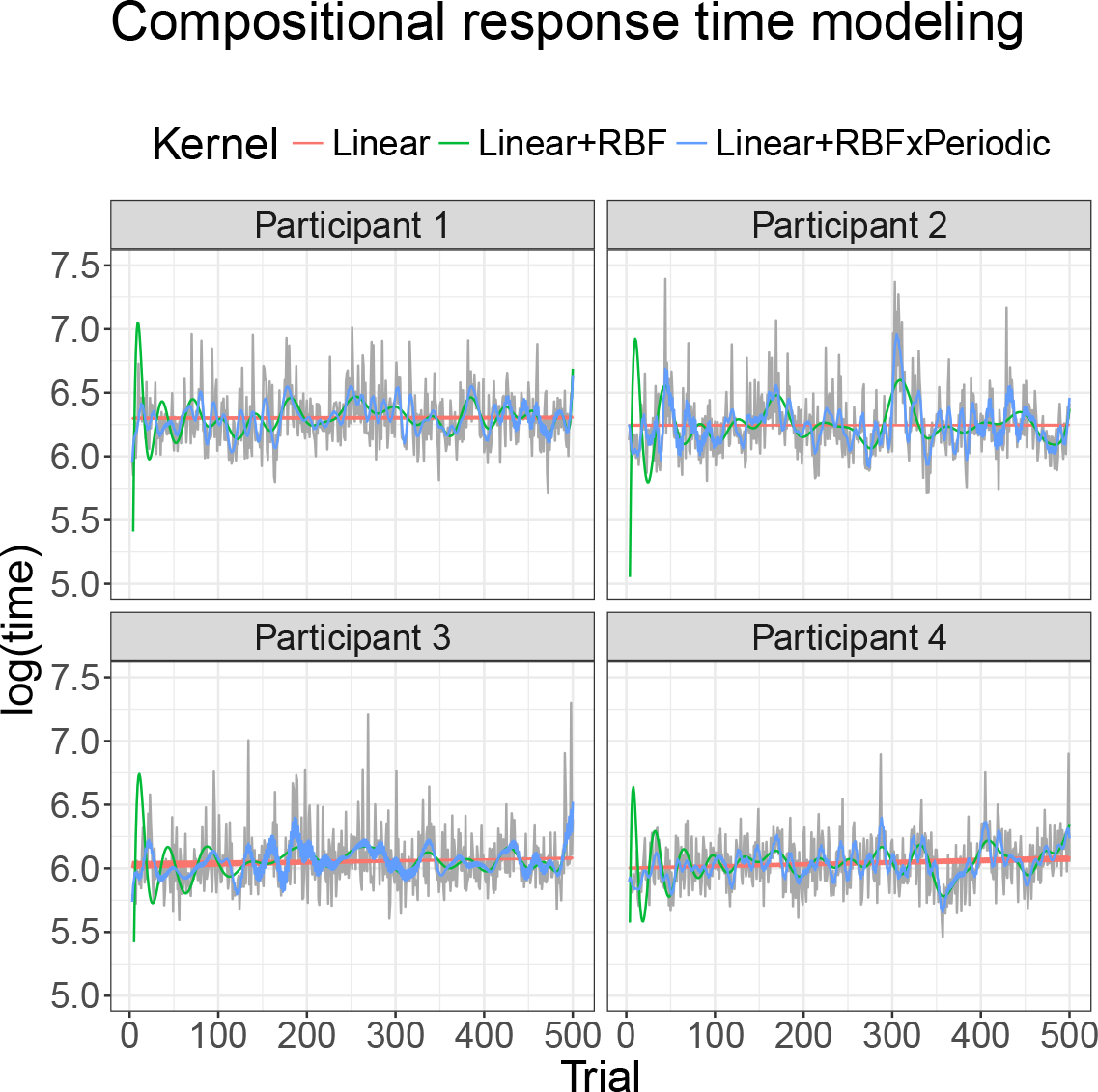
Response time data from Wagenmakers et al. Grey line shows raw log-response times. Coloured lines are created by the compositions extracted by compostional Gaussian Process regression.

Interestingly, all of the participants are best described by the same compositional components which are a Periodic × Linear + RBF (see Figure 7) which indicates a repeating pattern with increasing amplitude and an overall smooth inter-dependency between trials. This means that participants might be going through stages of shorter and longer response trials while the biggest effect is that trials are predicted by previous trials in a smooth way, similar to what has been found in the literature before.

## 5. General set-up for exploration-exploitation problems

Having found a powerful way to model functions, we can now focus on ways to cleverly explore and exploit unknown functions. Within the Gaussian process approach both pure *exploration* and *exploration-exploitation* can be treated in a similar manner. Both use Gaussian process regression to model the underlying function^5^ and estimate the utility of available queries (candidate input points to sample next) through what is called an *acquisition function.* An acquisition function *V* can be seen as measuring the usefulness (or utility) of candidate input points in terms of allowing one to learn the function as best as possible (exploration) or producing the best possible output (exploitation). The approach then goes on to choose as the next input the one that promises to produce the highest utility. The way this works is shown in Algorithm 1.

This algorithm starts out with a Gaussian process distribution over functions, then assesses the usefulness of the available samples by utilizing the acquisition function and selects the point that currently maximizes this function. The value of the utility function *V*_*t*_(**x**) thereby always depends on the current posterior of the Gaussian process at time point *t* (it can change on every trial). Afterwards, the new output at the chosen sample point is observed, the Gaussian process is updated, and the process starts anew. We will use a simple radial basis function kernel to model the unknown functions for all of the remaining examples. This choice is reasonable as in this setting, we need to choose an input from a bounded range of possible input points. As we do not have to extrapolate beyond the lower and upper bound of this range, modelling the function mostly consists of interpolation.

### Algorithm 1

General 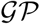 optimization algorithm

**Require**: Input space 𝓧 acquisition function V_t_; 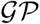-prior for *f* with mean
function *m*(x) and kernel *k*(x, x')
**for** t = 1, 2, … **do**
**Choose** 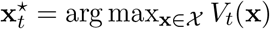
**Sample** 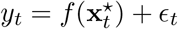
**end for**

## 6. Gaussian process active learning

The goal in a active learning setting is to learn an unknown function as accurately and quickly as possible. In a psychological setting this could mean for example to try and find out what a participant-specific forgetting curve might look like and choosing retention intervals adaptively in order to optimally learn about this function on each subsequent trial of an experiment (e.g., Myung, Cavagnaro, and Pitt, 2013).

### 6.1. *Acquisition function*

In Bayesian inference, learning about a function means that the posterior distribution over possible functions becomes more certain (e.g., less dispersed). A useful measure of the uncertainty about a random variable ***Y*** with probability distribution *p* is the differential entropy

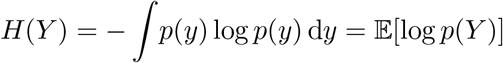

The information that an input *x* provides about the random variable, which we call the information gain, is the reduction in entropy due to observing the input and corresponding output

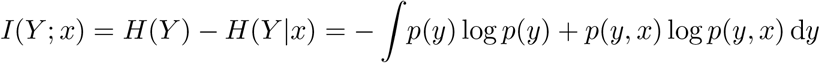

For example, if *Y* follows a *d*-variate Gaussian distribution with mean ***μ*** and covariance **Σ**, then the entropy is

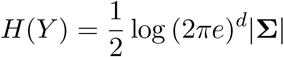

In our setting, we want to learn about the function, i.e. reduce the entropy in the distribution *p*(*f*). In Gaussian process regression, we can write the information gain as

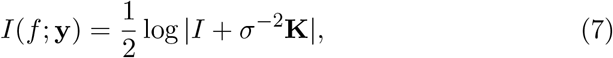

where K = [*k*(*x*,*x*′)].

Even though finding the overall information gain maximizer is NP-hard, it can be approximated by an efficient greedy algorithm based on Gaussian process regression. If *F*(*A*) = *I*(*f*; **y**_*A*_) is the information about the function *f* after having observed a set of points *A*, then this algorithm picks *x*_*t*_ = argmax *F*(*A*_*t*-1_ ⋃ {x}), that is greedily querying the point whose predicted output is currently most uncertain.

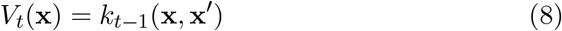

Here, uncertainty is measure by the variance of *f* at input **x**.

This algorithm starts with a Gaussian process prior for *f* and at each time *t* = 1,…,*T*, sequentially samples those input points where the current posterior predictive distribution 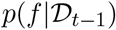 evaluated at **x** shows the highest variance, i.e. the highest predictive uncertainty. This is a “greedy” algorithm in the sense that it focuses on minimizing the current uncertainty, rather than looking further ahead into the future. Even though this algorithm, sometimes also called uncertainty sampling in statistics, looks na00EF;ve at first, it can actually be shown to obtain at least a constant fraction of the maximum information gain reachable using at most *T* samples (see Krause, Singh, and Guestrin, 2008, for more details):

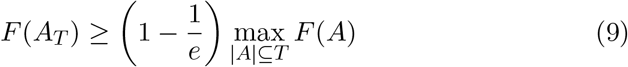

where *F*(*A_T_*) measures the information about *f* at time point *t* within the set *A* and *e* is Euler’s number. This is based on two properties of the acquisition function called *submodularity* and *monotonicity* (Krause and Golovin, 2012). Intuitively, submodularity here corresponds to a diminishing returns property of the acquisition function by which newly sampled points will add less and less information about the underlying function. Montonicity means that information never hurts (it is always helpful to observe more points). Both properties are crucial to show that the greedy algorithm can be successful. A simple example of the Gaussian process uncertainty reduction sampler is shown in Figure 8 below. We have used the same set of observations as before and let the algorithm select a new observation by picking as the next observation the one that currently has the highest predictive uncertainty attached.

**Figure 8:**
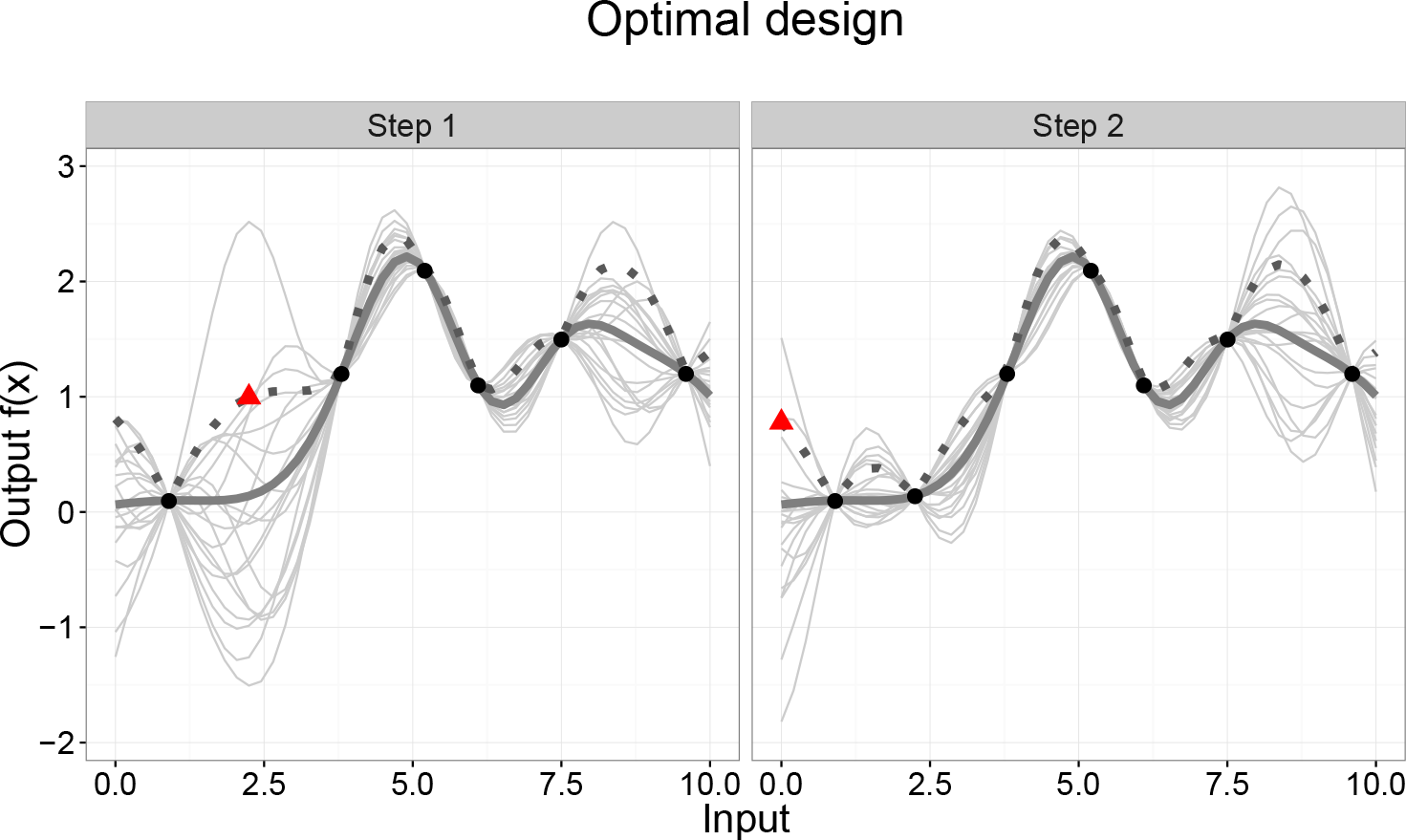
GP-uncertainty reduction example. The dark grey line marks the current mean of the GP. The dashed line shows the mean plus the standard deviation. The light grey lines are samples from the GP. The red triangle marks the current candidate point with the highest attached uncertainty.

### 6.2. *Example: Learning unknown functions*

In order to demonstrate how Gaussian process-based exploration works, we will show how the algorithm learns a set of unknown functions and compare it to other algorithms. The objective is to learn an unknown function as quickly and accurately as possible. For simplicity, we will focus on a function *f* which takes a one-dimensional and discretized input *x* ∈ [0,0.01,0.02,…, 10.00] and to which it maps an output *y*.

As GP regression is considered to learn many different functions well, we will test the algorithm on a number of different functions that are frequently encountered in psychology: a linear, quadratic, cubic, logarithmic, sine, and a non-stationary^6^ function. The functions are summarized in Table 4.

**Table 4.** Functions used in the Gaussian process exploration simulation.

| Function | Equation |
| --- | --- |
| linear | $f(x) = x$ |
| quadratic | $f(x) = x^2 + x$ |
| cubic | $f(x) = x^3 - x^2 + x$ |
| sine | $f(x) = x \times \sin(x)$ |
| non-stationary | $f(x) = \begin{cases} \sin(\pi x) + \cos(\pi x), & \text{if } x < 8 \\ x, & \text{otherwise} \end{cases}$ |

In addition to a GP regression model, we also used models that explicitly assume the function has a particular parametric form. These latter models learn the parameters (the weights) defining the function directly and were defined as a Gaussian process with a polynomial kernel with fixed degrees of freedom, i.e. performing Bayesian linear regression. All of the models were set up to learn the underlying function by picking as the next observation the one that currently has the highest uncertainty (standard deviation of the predicted mean) from within the input space *x* = [0,10].

We let each model run 100 times over 40 trials for each underlying function and averaged the mean squared error over the whole discretized input space for each step. We tested two different versions of learning the underlying functions with a Gaussian process regression, one which selected input points at random, i.e. uniformly from within the input space (GP-Random), and the uncertainty reduction sampler described above, which learns actively by choosing input points based on their predictive variance (GP-Active). For all models, on each trial, the hyper-parameters (e.g., the length-scale of the RBF kernel) were optimized by maximizing the marginal log likelihood of the observations thus far. Results are shown in Figure 9.

**Figure 9:**
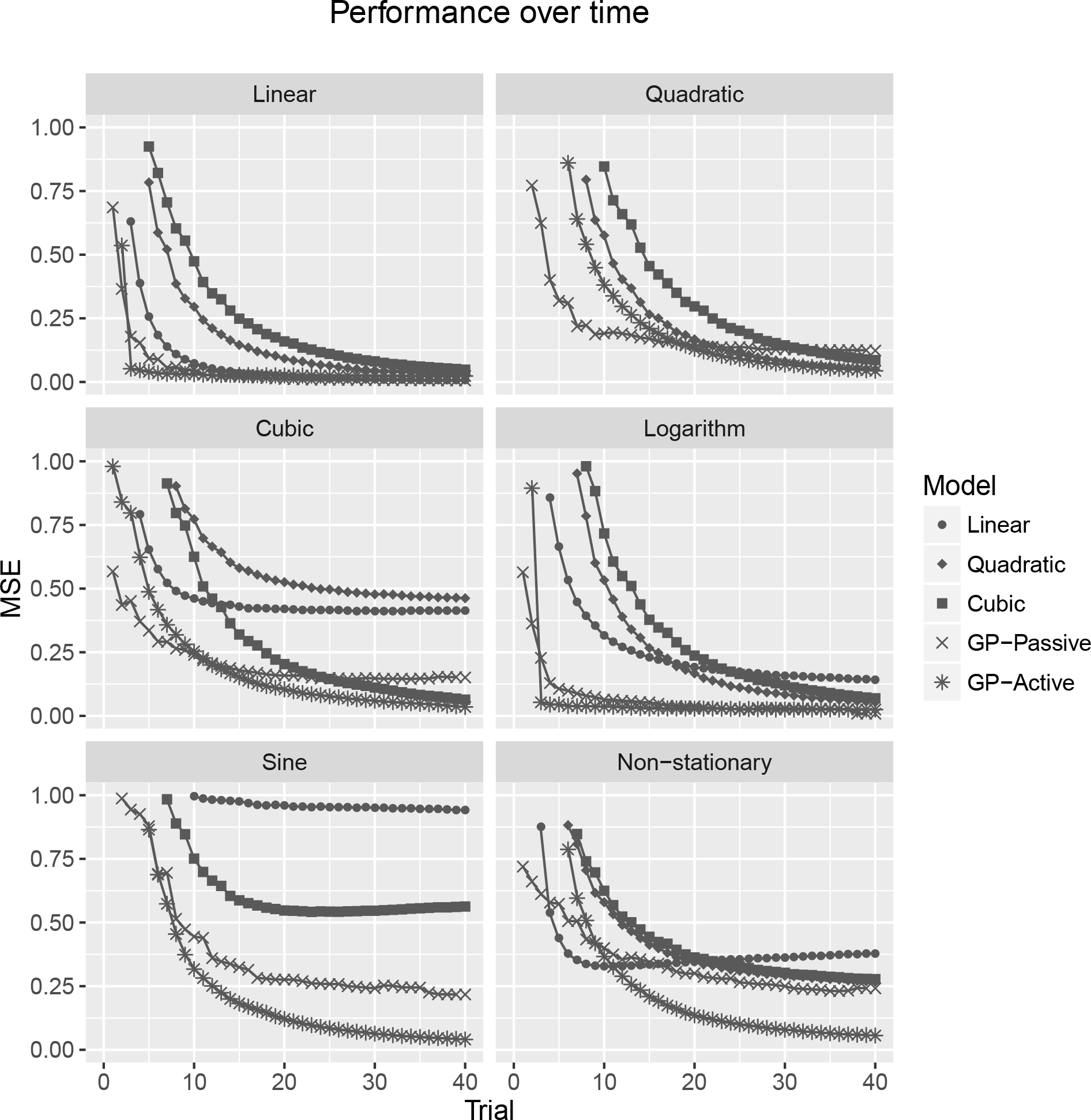
GP-uncertainty reduction example. GP-produced error always goes down. Linear model not always shown due to poor performance.

It can be seen that the Gaussian process model learns all functions efficiently. Even when the inputs are sampled at random, the error always goes down for a Gaussian process regression. However, the error generally goes down faster when inputs are selected actively. The other models only occasionally learn better than the GP models, when the assumed parametric form matches the true underlying form (for example, using a linear function to learn an underlying linear function). In some cases, using a cubic Bayesian regression seems to result in overfitting which leads to the overall error increasing again. In such cases, it might sometimes be better to select input points at random first. Overall, the results indicate that Gaussian process regression is especially useful in cases where the underlying function is not known.

## 7. Exploration-Exploitation and Bayesian Optimization

In an exploration-exploitation scenario the goal is to find the input to a function that produces the maximum output as quickly as possible.

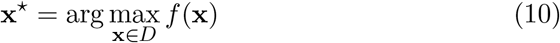

where **x**^⋆^ is the input that produces the highest output. One way to measure the quality of this search process is to quantify regret. Regret is the difference between the output of the currently chosen argument and the best output possible

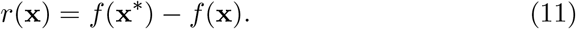

The cumulative regret is the sum of the regret over all trials, and the goal in an exploration-exploitation scenario is to minimize the cumulative regret:

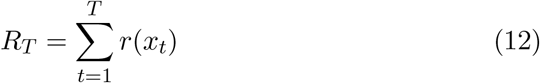

Again, finding the strategy that chooses the inputs to minimize the expected cumulative regret is NP-hard. That is, determining the sequence of queries (i.e. input choices) that lead to the lowest total regret is impossible for all but the most trivial cases. However, there is again a greedy trick one can apply in this scenario, which starts by reinterpreting the function maximization – or regret minimization – problem as a multi-armed bandit task (cf Katehakis and Veinott Jr, 1987). In a bandit task there are multiple options (arms) with unknown probability of producing a reward and the goal is to choose the best arm in order to maximise the overall reward (the name stems from the one armed bandits that can be found in casinos). In the current situation, we can view the discretized input points as the arms of a multi-armed bandit, and the output of the function at those points as the unknown rewards that are associated to each arm. What distinguishes the current situation from traditional bandit tasks is that the rewards of the arms are correlated in dependency of the underlying covariance kernel. Nevertheless, viewing the task as a multi-armed bandit allows us to use strategies that have been devised for traditional bandit tasks. One popular strategy is called the upper confidence bound (UCB) algorithm, which relies on the following acquisition function:

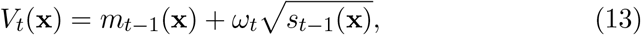

where 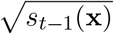 is the predictive standard deviation at a point **x**, and *m*_*t*_ is the posterior mean function (4) and the posterior variance is *s*_*t*_ = *k*_*t*_(**x**, **x**) (5). Finally, *ω*_*t*_ is a free parameter that determines the width of the confidence interval. For example, setting *ω*_*t*_ = 1.96, results in a 95% confidence interval for a single value **x** given a Gaussian distribution.

The UCB algorithm chooses the arm for which the upper confidence bound is currently the highest. The upper confidence bound is determined by two factors: the current estimate of the mean of *f* at a particular point (the higher the estimate, the higher the bound) and the uncertainty attached to that estimate (the higher the uncertainty, the higher the bound). Therefore, the UCB algorithm trades off naturally between expectation and uncertainty. An example of how the UCB algorithm works, using the same data as before, is shown in Figure 10.

**Figure 10:**
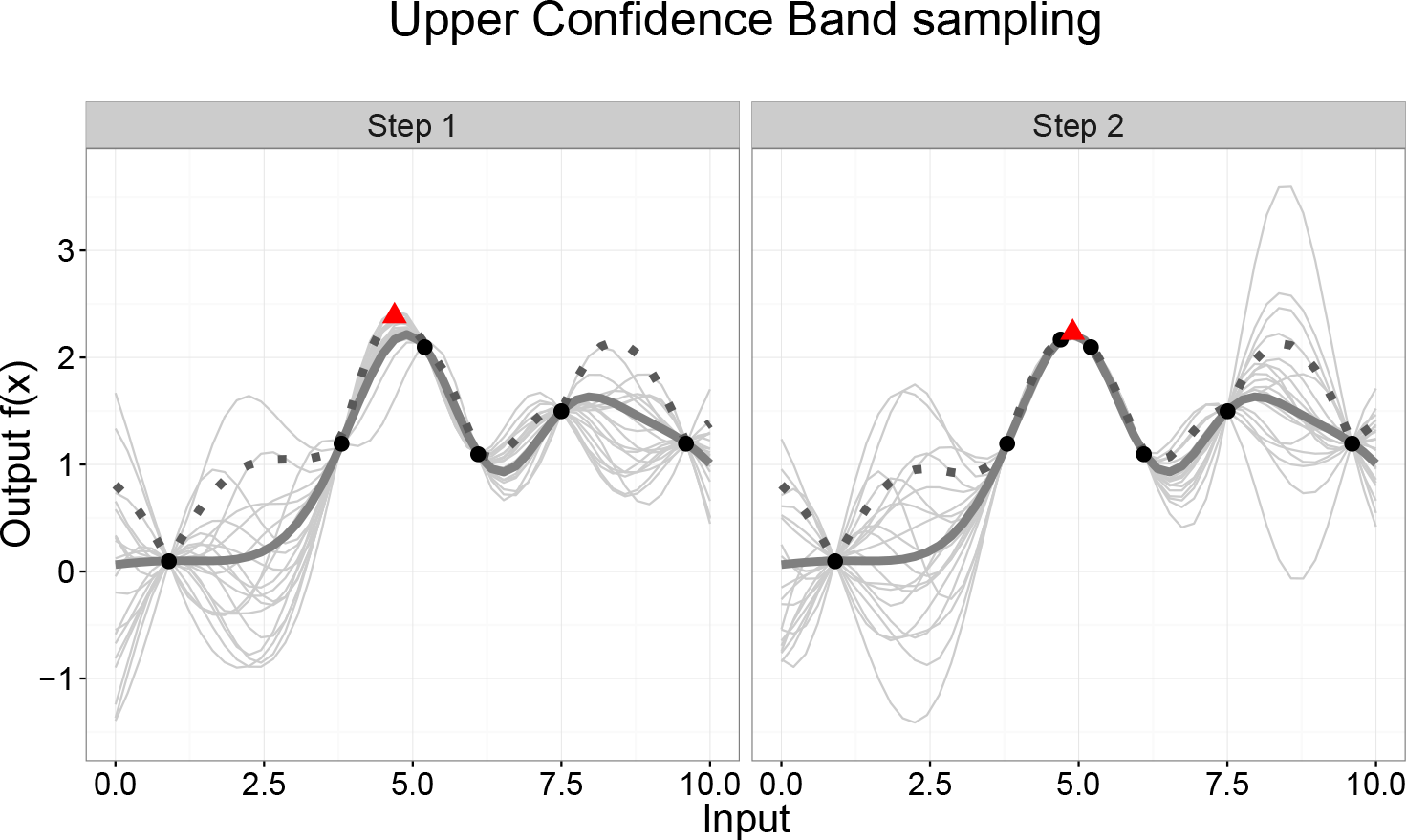
GP-UCB example. The dark grey line marks the current mean of the GP. The dashed line marks the GP’s upper confidence bound. The light grey lines are samples from the GP. The red triangle marks the point that currently produces the highest UCB.

Even though the greedy UCB strategy is naïve, it can be shown that its regret is sublinear for suitable choices of *ω*_*t*_, again using an argument that relies on the submodularity and monotonicity of the overall information gain (Srinivas, Krause, Kakade, and Seeger, 2009). Sublinear regret here just means that the regret per round goes down in expectation, thereby guaranteeing that the algorithm picks better points over time. These regret bounds are known even for the agnostic case in which the underlying function is unknown (but lies in the RKHS norm, see Srinivas et al., 2009). However, trying to optimize an underlying function with the wrong prior kernel can lead to a noticeable increase in regret (Schulz et al., 2016c).

### 7.1. *GP-UCB Example: Recommending movies*

As an example of applying Gaussian Process upper confidence bound sampling (GP-UCB) to exploration-exploitation problems, we will use it in a movie recommendation scenario, where the task is to recommend the best movies possible to a user with unknown preferences. This involves both learning how different features of movies influence the liking of a movie and recommending the movies that will be liked the most. For this application, we sampled 5141 movies from the IMDb database and recorded their features such as the year they appeared, the budget that was used to make them, their length, as well as how many people had evaluated the movie on the platform, number of facebook likes of different actors within the movie, genre of the movie, etc. As a proxy for how much a person would enjoy the movie, we used the average IMDb score, which is based on the ratings of registered users. As there were 27 features in total, we performed a Principal Component Analysis extracting 8 components that together explained more than 90% of the variance within the feature sets. These components were then used as an input for the optimization routine. We used a GP-UCB with a radial basis function kernel, set *ω* = 3 in the UCB acquisition function to encourage exploration ^7^, initialized the GP with 5 randomly sampled observations, and then let the algorithm pick 20 movies sequentially. This procedure was repeated 50 times. Even though recommender systems normally try to recommend the best movie for a particular user, this approach can be seen as recommending movies to an average user.

**Figure 11:**
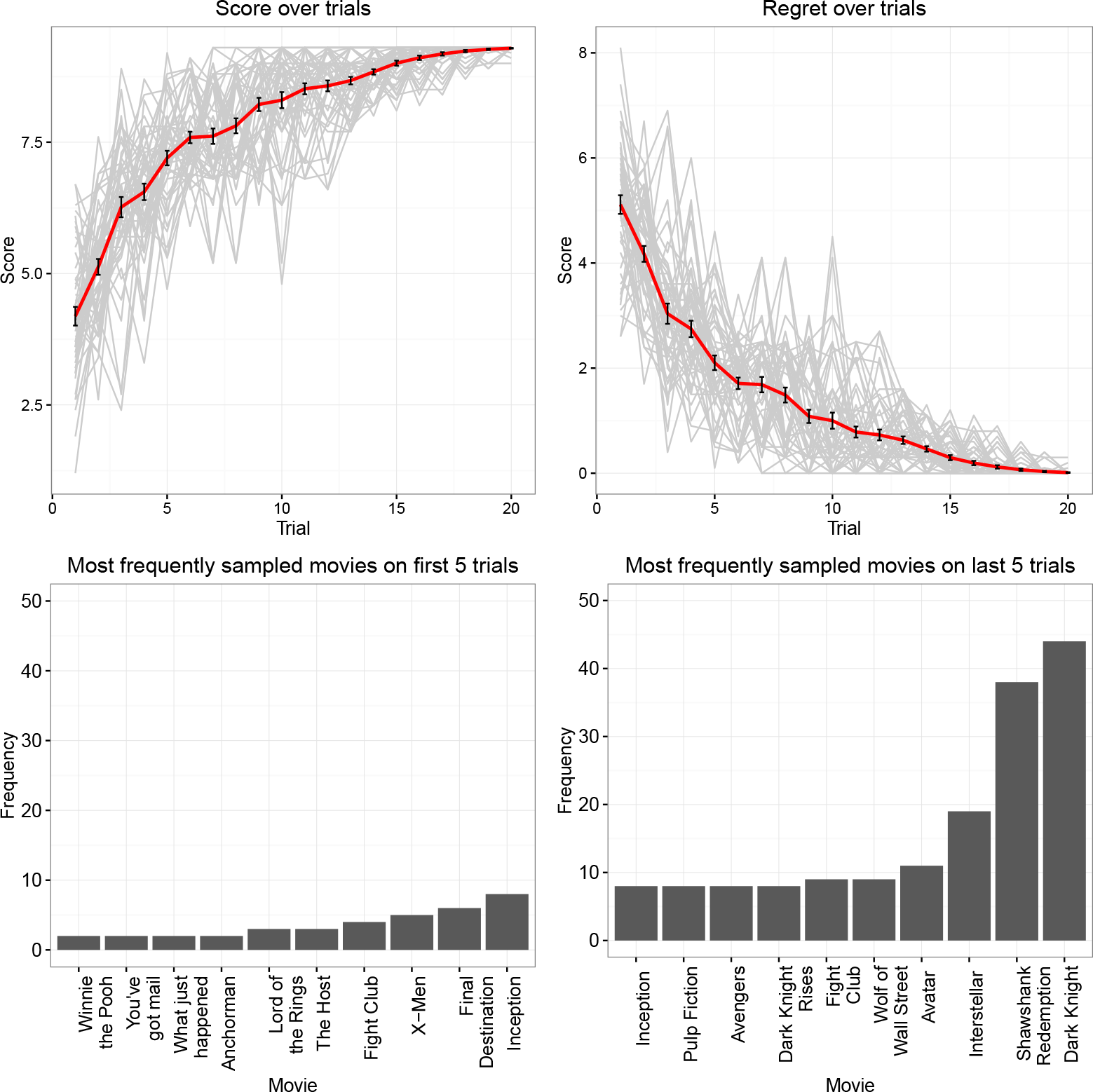
Recommending movies with a GP-UCB algorithm. The score (upper left, error bars represent the standard error of the mean) goes up over all runs and plateaus very quickly at around the highest value possible (9.3). Vice versa, the overall regret (upper right) goes down over trials an quickly approaches 0. Within the first 5 samples, movies are mostly picked at random and no clear pattern of movies seems to dominate (bottom right). However, within the last 5 trials GP-UCB preferentially samples highly rated movies (bottom right).

Results are shown in Figure 11. It can be seen that the algorithm quickly starts choosing movies that produce high scores which results in the overall mean score to go up and the regret to go down over time. Moreover, the variance of the picked movies also goes down over time as GP-UCB almost exclusively samples highly rated movies later on. Whereas the 10 most frequently sampled movies within the first 5 samples seem to be sampled closely to random, the most frequently sampled movies within the last 5 trials are movies that are generally highly rated. In the end, the algorithm has explored the input space well, learned the unknown preference function of the average user rather well, and returned movies that are on average highly rated. When we let the GP-UCB algorithm run over 200 trials, it frequently starts sampling the movie “The Shawshank Redemption”, which is the highest rated movie on the internet movie database.

## 8. Safe exploration-exploitation

Sometimes an exploration-exploitation scenario may come with additional requirements. For example, one such requirement can be to avoid certain outputs. Consider excitatory stimulation treatment, where the task is to stimulate the spinal chord in such a way that certain movements are achieved (Desautels, Choe, Gad, Nandra, Roy, Zhong, Tai, Edgerton, and Burdick, 2015). Here, it is important to stimulate the spinal chord such that optimal recovery is obtained, but not too much as this might lead to painful reactions for the patients.

Again, Gaussian process optimization methods can be used to learn the underlying function of what stimulation leads to which strength of reaction. However, an additional requirement now is to avoid particularly reactions that result in pain. An algorithm that balances exploration and exploitation whilst avoiding certain outputs is called Safe Optimization (Sui, Gotovos, Burdick, and Krause, 2015). This algorithm adapts the Upper Confidence Bound approach described earlier to accommodate this additional requirement. It works by trading-off two different goals: Firstly, it keeps track of a set of safe options it considers to be above a given safe threshold (points currently showing a high likelihood of being above the threshold) and tries to expand this set as much as it can. Secondly, it maintains a set of potential maximizers (points likely to produce high outcomes) that, if used as an input, would potentially achieve the highest output. It then chooses as the next input a point within the intersection of these two sets, that is a safe point that is either a maximizer or an expander that has the highest predictive variance and potentially expands the set of maximisers. This algorithm can also be adapted to separate the objective function from a set of constraints as described by (Berkenkamp, Krause, and Schoellig, 2016).

More formally, a *safe set* of possible inputs that are likely to provide outputs above the threshold is defined and then further separated into a set of *maximizers* (inputs that promise to provide the maximum output) and *expanders* (inputs that promise to expand the safe set). This algorithm uses the upper and lower bounds of a confidence interval as described in Equation 13 above, i.e. by either setting *ω* to 3 or −3 for the upper and lower confidence bound respectively. Using these bounds, it is possible to define the safe set as all the input points in the set of available inputs whose lower confidence bound is above the provided threshold. This is intuitive as one would expect these points to be above the threshold in 0.1% of the cases. The set of potential maximizers contains all safe inputs that promise to obtain the maximum output value; these are the safe inputs for which their upper confidence bound is above the highest lower bound within the input set, i.e. points with an upper bound better than the best lower bound. The set of expanders is normally found by forward simulations, where it is assessed if the safe set is –in expectation– expanded by sampling a given point. For further technical details, we refer the interested reader to Sui et al. (2015).

### 8.1. *Example: Cautious stimulus optimization*

As an illustration of the Safe Optimization algorithm, we apply it to a situation in which the objective is to choose inputs x in order to learn about the underlying function in a two-dimensional space such that –eventually– points that produce high outputs in *y* will be sampled whilst avoiding to choose inputs that produce an output below 0. To simplify presentation, we sampled the underlying function from a Gaussian process parameterized by a radial basis function kernel. This can be seen as similar to the case where one wants to present stimuli to participants, but make sure that participants never react with an intensity below a certain threshold.

Results are shown in Figure 12. It can be seen that the Safe Optimization algorithm explores the function exceptionally well in its attempt to expand the space of possible safe inputs. At the same time, the algorithm does not at any time choose inputs from the white area (producing output values below 0). This algorithm could be applied to optimal design settings that require additional constraints.

**Figure 12:**
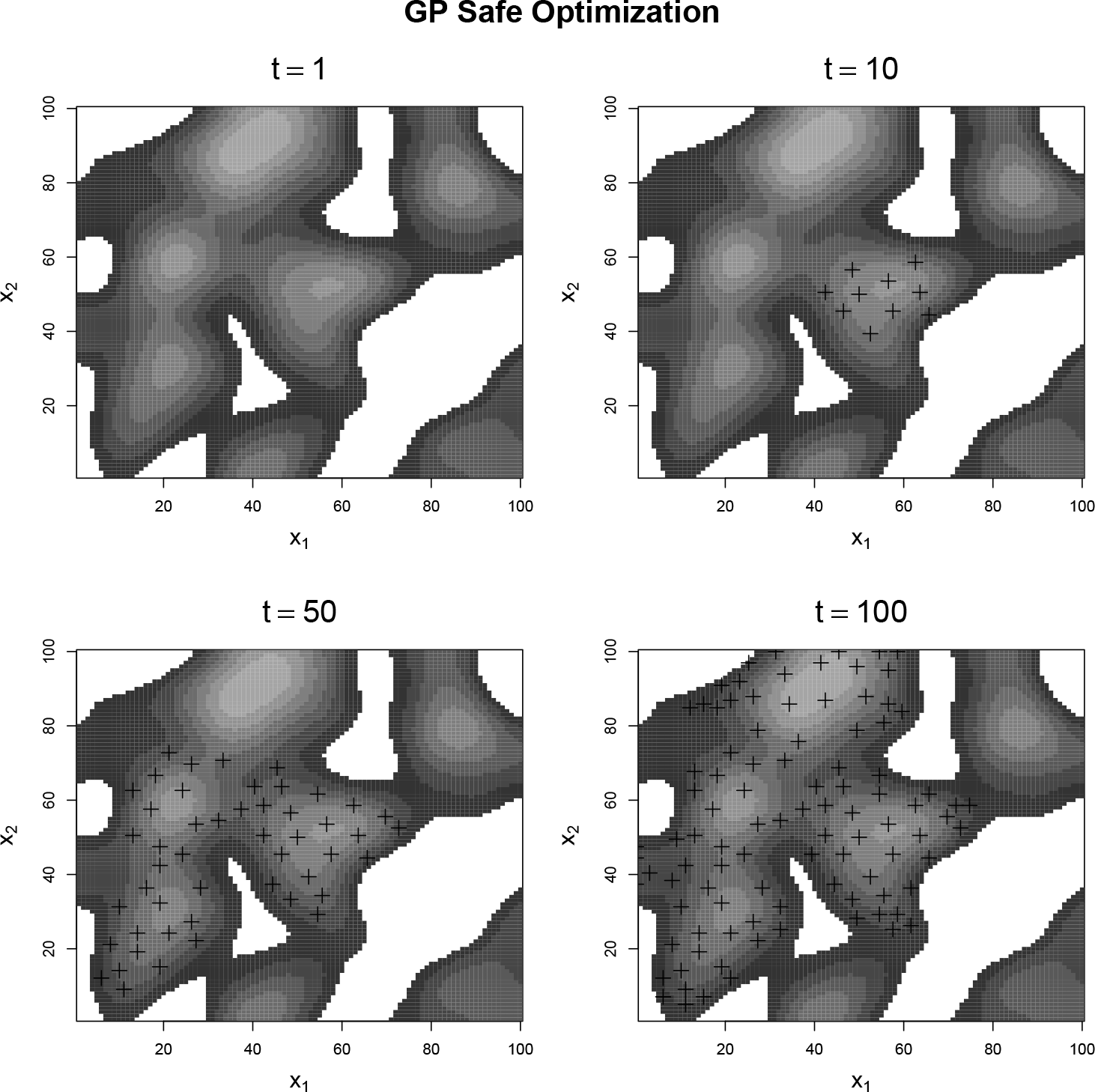
GP-Safe Optimization example showing samples after 1,10, 50 and 100 samples. White represents areas below 0. The black crosses show where the Safe Optimization algorithm has sampled. Lighter areas represent higher scores. The algorithm efficiently explores other safe areas. It never samples points within the surrounding white area as these are below the threshold.

## 9. Gaussian processes and cognition

We have seen that Gaussian process regression is a powerful tool to model, explore, and exploit unknown functions. However, Gaussian process regression might also be applied in a different, more psychological context, namely as a model of human cognition in general and function learning in particular. Recently, Lucas, Griffiths, Williams, and Kalish (2015) have proposed to use Gaussian process regression as a rational model of function learning that can explain various effects within the literature of human function learning. Schulz, Tenenbaum, Reshef, Speekenbrink, and Gershman (2015) used Gaussian processes to assess participants’ judgements of the predictability of functions in dependency of the smoothness of the underlying kernel. As many different kernels can be used to model function learning, Wilson, Dann, Lucas, and Xing (2015) tried to infer backwards what the human kernel might look like by using a non-parametric kernel approach to Gaussian process regression. As explained above, kernels can also be added together and multiplied to build more expressive kernels, which led Schulz, Tenenbaum, Duvenaud, Speekenbrink, and Gershman (2016d) to assess if participants’ functional inductive biases can be described as made up of compositional building blocks. In a slightly different context, Gershman, Malmaud, Tenenbaum, and Gershman (2016) modelled participants– utility of combinations of different objects by a Gaussian process parametrized by a tree-like kernel.

Within an exploration-exploitation context, Borji and Itti (2013) and Wu, Schulz, Speekenbrink, Nelson, and Meder (2017) showed that Gaussian process-based optimization can explain how participants actively search for the best output when trying to optimize one-dimensional functions.Schulz, Konstantinidis, and Speekenbrink (2016b) used Gaussian process exploration-exploitation algorithms to model behaviour in tasks that combine function learning and decision making (contextual multi-armed bandit tasks). Lastly, Schulz, Huys, Bach, Speekenbrink, and Krause (2016a) applied the safe optimization algorithm described here to scenarios in which participants had to cautiously optimize functions while never sampling below a given threshold.

## 10. Discussion

This tutorial has introduced Gaussian process regression as a general purpose technique to model, explore and exploit unknown functions. We have mainly focused on Gaussian process regression with a radial basis function kernel, but many other kernels and kernel combinations are possible and –as we have indicated above– many standard Bayesian regression approaches can be re-parametrized to be equivalent to Gaussian process regression, given specific assumptions about the kernel (Duvenaud et al., 2013).

Of course a tutorial like this can never be fully comprehensive. For example, many other acquisition functions than the ones introduced here (uncertainty sampling and UCB) exist. For pure exploration, another commonly used acquisition function attempts to minimize the expected variance over the whole input space (Gramacy and Apley, 2014). This method tends to sample less on the bounds of the input space, but can be hard to compute, especially if the input space is large. There also exist many different acquisition functions in the exploration-exploitation context, that are mostly discussed under the umbrella term Bayesian optimization (de Freitas, Smola, and Zoghi, 2012). Two other acquisition functions that are frequently applied here are the *probability of improvement* and the *expected improvement* (Močkus, 1975), which choose inputs that have a high probability to produce a better output than the input that is currently estimated to be best, or that produce an output which is expected to surpass the expected outcome of the input currently thought best. Thompson sampling (Thompson, 1933; May, Korda, Lee, and Leslie, 2012) is another acquisition function, which chooses an action that maximizes the expected outcome with respect to a randomly drawn belief, and has recently gained popularity because of its competitive empirical performance (Chapelle and Li, 2011).

Another situation in which Gaussian processes are frequently applied is called “global optimization”, in which the goal is finding the overall maximum of a function as quickly as possible, but without worrying about the outputs that were produced in the search process. Parameter estimation is an example of such a problem and again different algorithms have been proposed, in particular the proposal by Hennig and Schuler (2012) to maximize the information gain about the location of the maximum. There is also a growing community of researchers who apply Gaussian process-based algorithms to return uncertainty estimates of traditional computational methods such as optimization, quadrature, or solving differential equations under the umbrella term “probabilistic numerics” (Hennig, Osborne, and Girolami, 2015).

Gaussian process regression does have some drawbacks. One such drawback, as compared to traditional regression models, is that parameter-based interpretations such as “if *x* increases by 1, *y* increases by 2” are not directly possible. However, as different kernels encode different assumptions about the underlying functions, assessing which kernel describes the underlying function best can be used as a basis to interpret the modelled function (Lloyd et al., 2014). Choosing the appropriate kernel is a difficult problem. General solutions to this are to construct more complicated kernels from a set of relatively simple base kernels (as shown above) and to search the kernel space by proposing and checking new kernel combinations (Duvenaud et al., 2013), or to define the kernel in a non-parametric manner by using a non-parametric approach towards estimating the kernel itself (Wilson and Adams, 2013). Possibly the biggest drawback of Gaussian process regression is its poor scaling. As inferring the posterior involves inverting the matrix 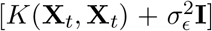, inference scales cubically with the number of observations^8^. Speeding up inference for Gaussian process regression therefore is a topic of ongoing research. Some methods that have been proposed are to sparsely approximate inputs (Lawrence, Seeger, and Herbrich, 2003) or to bound the computational cost of the matrix inversion by projecting into a pre-defined finite basis of functions drawn from the eigen-spectrum of the kernel (Rahimi and Recht, 2007).

We hope to have shown some interesting examples of Gaussian process regression as a powerful tool for many applied situations, specifically exploration-exploitation scenarios, and hope that this tutorial will inspire more scientists to apply these methods in the near future. Currently available software that can assist in this is listed in Table 5.

**Table 5.**
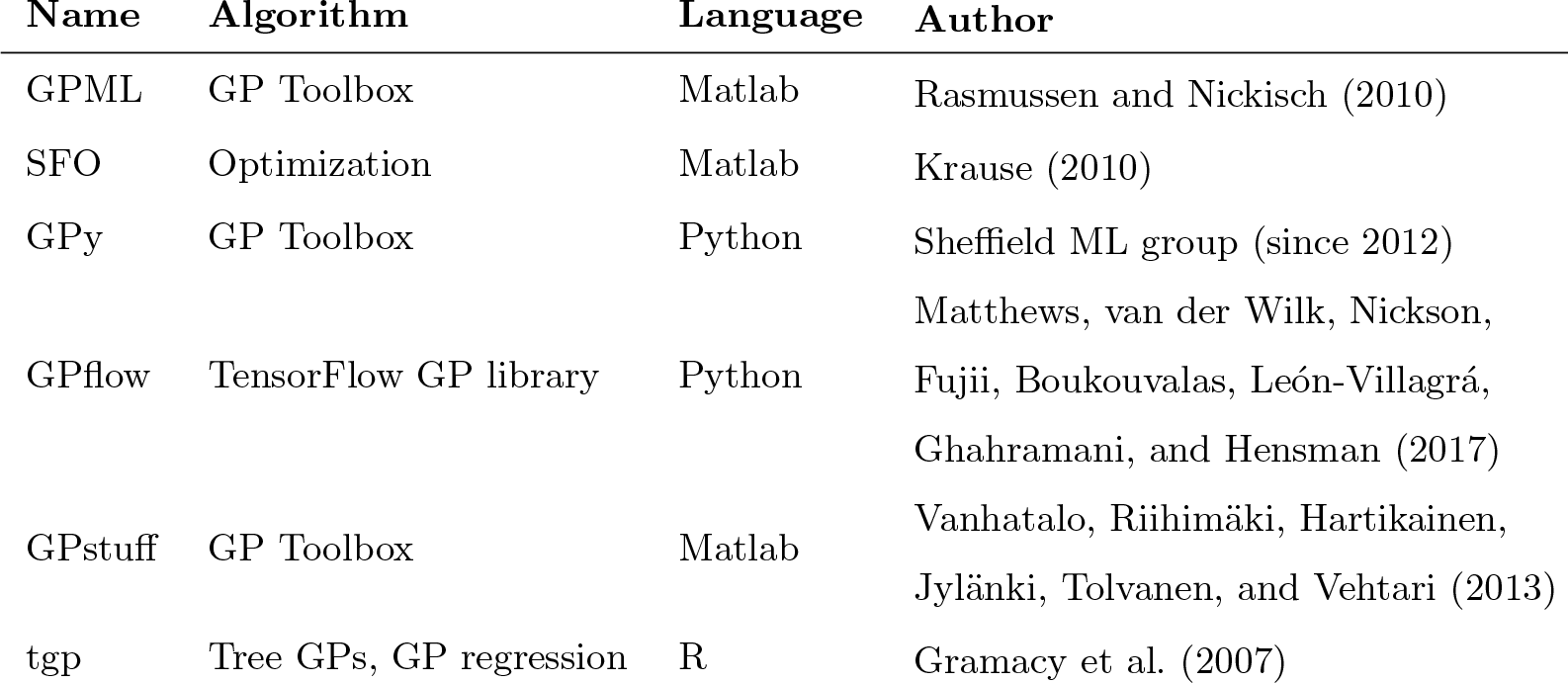
Gaussian process packages

## Footnotes

1 We will see later that it is in fact not only the data that determines the complexity of the Gaussian process, but also the chosen kernel.

2 In fact, simple Bayesian linear regression can be recovered by using a linear kernel 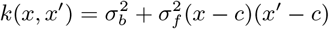, which means that for 0-mean, *k*(*x*; *x*′) = *x*^T^*x*′

3 This is also sometimes referred to as the “kernel trick”

4 A recent version of this function is available at http://learning.eng.cam.ac.uk/carl/code/minimize

5 In this context, a Gaussian process regression is sometimes also referred to as a “surrogate model” (see Gramacy and Lee, 2008).

6 A non-stationary function for our purpose is a function that changes its parametric form over different parts of the input space

7 Running the algorithm with *ω* = 2 or setting *ω* dynamically leads to similar results.

8 This is the computational complexity; the regret remains the same as before.

## References

Akaike, H., 1974. A new look at the statistical model identification. IEEE transactions on automatic control 19 (6), 716–723.

Berkenkamp, F., Krause, A., Schoellig, A. P., 2016. Bayesian optimization with safety constraints: safe and automatic parameter tuning in robotics. arXiv preprint arXiv:1602.04450.

Borji, A., Itti, L., 2013. Bayesian optimization explains human active search. In: Advances in Neural Information Processing Systems. pp. 55–63.

Cavagnaro, D. R., Aranovich, G. J., McClure, S. M., Pitt, M. A., Myung, J. I., 2014. On the functional form of temporal discounting: An optimized adaptive test.

Chapelle, O., Li, L., 2011. An empirical evaluation of thompson sampling. In: Advances in neural information processing systems. pp. 2249–2257.

Cox, G., Kachergis, G., Shiffrin, R., 2012. Gaussian process regression for trajectory analysis. In: Proceedings of the Cognitive Science Society. Vol. 34.

de Freitas, N., Smola, A., Zoghi, M., 2012. Regret Bounds for Deterministic Gaussian Process Bandits. arXiv preprint arXiv:1203.2177.

Desautels, T. A., Choe, J., Gad, P., Nandra, M. S., Roy, R. R., Zhong, H., Tai, Y.-C., Edgerton, V. R., Burdick, J. W., 2015. An active learning algorithm for control of epidural electrostimulation. IEEE Transactions on Biomedical Engineering 62 (10), 2443–2455.

Durrleman, S., Simon, R., 1989. Flexible regression models with cubic splines. Statistics in medicine 8 (5), 551–561.

Duvenaud, D., Lloyd, J. R., Grosse, R., Tenenbaum, J. B., Ghahramani, Z., 2013. Structure discovery in nonparametric regression through compositional kernel search. arXiv preprint arXiv:1302.4922.

Engbert, R., Kliegl, R., 2004. Microsaccades keep the eyes’ balance during fixation. Psychological science 15 (6), 431–431.

Flaxman, S., Gelman, A., Neill, D., Smola, A., Vehtari, A., Wilson, A. G., 2015. Fast hierarchical gaussian processes.

Freeman, J. B., Ambady, N., 2010. Mousetracker: Software for studying real-time mental processing using a computer mouse-tracking method. Behavior Research Methods 42 (1), 226–241.

Gershman, S. J., Blei, D. M., 2012. A tutorial on Bayesian nonparametric models. Journal of Mathematical Psychology 56 (1), 1–12.

Gershman, S. J., Malmaud, J., Tenenbaum, J. B., Gershman, S., 2016. Structured representations of utility in combinatorial domains.

Gramacy, R. B., Apley, D. W., 2014. Local gaussian process approximation for large computer experiments. Journal of Computational and Graphical Statistics (just-accepted), 1–28.

Gramacy, R. B., Lee, H. K., 2008. Bayesian treed Gaussian process models with an application to computer modeling. Journal of the American Statistical Association 103 (483).

Gramacy, R. B., et al., 2007. tgp: an R package for Bayesian nonstationary, semiparametric nonlinear regression and design by treed Gaussian process models.

Hennig, P., Osborne, M. A., Girolami, M., 2015. Probabilistic numerics and uncertainty in computations. Proc. R. Soc. A 471 (2179), 20150142.

Hennig, P., Schuler, C. J., 2012. Entropy search for information-efficient global optimization. Journal of Machine Learning Research 13 (Jun), 1809–1837.

Jäkel, F., Schölkopf, B., Wichmann, F. A., 2007. A tutorial on kernel methods for categorization. Journal of Mathematical Psychology 51 (6), 343–358.

Kac, M., Siegert, A., 1947. An explicit representation of a stationary gaussian process. The Annals of Mathematical Statistics, 438–442.

Katehakis, M. N., Veinott Jr, A. F., 1987. The multi-armed bandit problem: decomposition and computation. Mathematics of Operations Research 12 (2), 262–268.

Kieslich, P. J., Henniger, F., 2017. Mousetrap: An integrated, open-source mouse-tracking package. Behavioral Research Methods.

Krause, A., 2010. Sfo: A toolbox for submodular function optimization. Journal of Machine Learning Research 11 (Mar), 1141–1144.

Krause, A., Golovin, D., 2012. Submodular function maximization. Tractability: Practical Approaches to Hard Problems 3, 19.

Krause, A., Singh, A., Guestrin, C., 2008. Near-optimal sensor placements in gaussian processes: Theory, efficient algorithms and empirical studies. The Journal of Machine Learning Research 9, 235–284.

Lawrence, N., Seeger, M., Herbrich, R., 2003. Fast sparse gaussian process methods: The informative vector machine. In: Proceedings of the 16th Annual Conference on Neural Information Processing Systems. No. EPFL-CONF-161319. pp. 609–616.

Lee, C. H., 2004. A phase space spline smoother for fitting trajectories. IEEE Transactions on Systems, Man, and Cybernetics, Part B (Cybernetics) 34 (1), 346–356.

Lloyd, J. R., Duvenaud, D., Grosse, R., Tenenbaum, J. B., Ghahramani, Z.,2014. Automatic construction and natural-language description of non-parametric regression models. arXiv preprint arXiv:1402.4304.

Lucas, C. G., Griffiths, T. L., Williams, J. J., Kalish, M. L., 2015. A rational model of function learning. Psychonomic bulletin & review, 1–23.

Matthews, A. G. d. G., van der Wilk, M., Nickson, T., Fujii, K., Boukouvalas, A., León-Villagrá, P., Ghahramani, Z., Hensman, J., 2017. Gpflow:A gaussian process library using tensorflow. Journal of Machine Learning Research 18 (40), 1–6.

May, B. C., Korda, N., Lee, A., Leslie, D. S., 2012. Optimistic bayesian sampling in contextual-bandit problems. Journal of Machine Learning Research 13 (Jun), 2069–2106.

Meder, B., Nelson, J. D., 2012. Information search with situation-specific reward functions. Judgment and Decision Making 7 (2), 119–148.

Močkus, J., 1975. On Bayesian methods for seeking the extremum. In: Optimization Techniques IFIP Technical Conference. Springer, pp. 400–404.

Myung, J. I., Cavagnaro, D. R., Pitt, M. A., 2013. A tutorial on adaptive design optimization. Journal of mathematical psychology 57 (3), 53–67.

Myung, J. I., Pitt, M. A., 2009. Optimal experimental design for model discrimination. Psychological review 116 (3), 499.

Rahimi, A., Recht, B., 2007. Random features for large-scale kernel machines. In: Advances in neural information processing systems. pp. 1177–1184.

Rasmussen, C. E., Nickisch, H., 2010. Gaussian processes for machine learning (gpml) toolbox. Journal of Machine Learning Research 11 (Nov), 3011–3015.

Schulz, E., Huys, Q. J., Bach, D. R., Speekenbrink, M., Krause, A., 2016a. Better safe than sorry: Risky function exploitation through safe optimization. arXiv preprint arXiv:1602.01052.

Schulz, E., Konstantinidis, E., Speekenbrink, M., 2016b. Putting bandits into context: How function learning supports decision making. Journal of Experimental Psychology: Learning, Memory, and Cognition.

Schulz, E., Speekenbrink, M., Hernández-Lobato, J. M., Ghahramani, Z., Gershman, S. J., 2016c. Quantifying mismatch in bayesian optimization. In: Nips workshop on bayesian optimization: Black-box optimization and beyond.

Schulz, E., Tenenbaum, J. B., Duvenaud, D., Speekenbrink, M., Gershman, S. J., 2016d. Probing the compositionality of intuitive functions. Tech. rep., Center for Brains, Minds and Machines (CBMM).

Schulz, E., Tenenbaum, J. B., Reshef, D. N., Speekenbrink, M., Gershman, S. J., 2015. Assessing the perceived predictability of functions. In: Proceedings of the Thirty-Seventh Annual Conference of the Cognitive Science Society.

Sheffield ML group, since 2012. GPy: A gaussian process framework in python. http://github.com/SheffieldML/GPy.

Srinivas, N., Krause, A., Kakade, S. M., Seeger, M., 2009. Gaussian process optimization in the bandit setting: No regret and experimental design. arXiv preprint arXiv:0912.3995.

Sui, Y., Gotovos, A., Burdick, J., Krause, A., 2015. Safe exploration for optimization with gaussian processes. In: Proceedings of the 32nd International Conference on Machine Learning (ICML-15). pp. 997–1005.

Thompson, W. R., 1933. On the likelihood that one unknown probability exceeds another in view of the evidence of two samples. Biometrika 25 (3/4), 285–294.

Van Zandt, T., Townsend, J. T., 2014. Designs for and analyses of response time experiments. The Oxford Handbook of Quantitative Methods: Foundations 1, 260.

Vanhatalo, J., Riihimäki, J., Hartikainen, J., Jylänki, P., Tolvanen, V., Vehtari, A., 2013. Gpstuff: Bayesian modeling with gaussian processes. Journal of Machine Learning Research 14 (Apr), 1175–1179.

Wagenmakers, E.-J., Farrell, S., Ratcliff, R., 2004. Estimation and interpretation of 1/fα noise in human cognition. Psychonomic bulletin & review 11 (4), 579–615.

Wetzels, R., Vandekerckhove, J., Tuerlinckx, F., Wagenmakers, E.-J., 2010. Bayesian parameter estimation in the expectancy valence model of the iowa gambling task. Journal of Mathematical Psychology 54 (1), 14–27.

Williams, C. K., 1998. Prediction with gaussian processes: From linear regression to linear prediction and beyond. In: Learning in graphical models. Springer, pp. 599–621.

Williams, C. K., Rasmussen, C. E., 2006. Gaussian processes for machine learning. the MIT Press 2 (3), 4.

Wilson, A. G., Adams, R. P., 2013. Gaussian process kernels for pattern discovery and extrapolation. In: ICML (3). pp. 1067–1075.

Wilson, A. G., Dann, C., Lucas, C., Xing, E. P., 2015. The human kernel. In: Advances in Neural Information Processing Systems. pp. 2854–2862.

Wu, C. M., Schulz, E., Speekenbrink, M., Nelson, J. D., Meder, B., 2017. Exploration and generalization in vast spaces. bioRxiv, 171371.

